# Derivation of functional retinal endothelial cells from human pluripotent stem cells for therapeutics and modeling

**DOI:** 10.1101/2025.03.04.641453

**Authors:** Ying-Yu Lin, Parker Esswein, Lucas Ramirez, Emily Warren, Sharon Gerecht

**Affiliations:** Department of Biomedical Engineering, Duke University, Durham, NC, 27708, USA

## Abstract

Retinal microvascular diseases involve a compromised inner blood-retina barrier (iBRB), which remains poorly understood. A renewable source of human iBRB endothelium is thus vital for advancing eye research and treatment development. Here, we differentiated human iPSCs into retinal endothelial cells (iRECs) via the Wnt/β-catenin pathway, namely Norrin/Frizzled4 signaling. These iRECs show genetic, protein, and functional fidelity and unique retinal features. When injected into oxygen-induced retinopathy mice, iRECs integrated into the host vascular network and revascularized the ischemic eye, rescuing the tissue. Within microphysiological models, iRECs form perfusable microvascular networks that mimic the iBRB’s morphology and phenotype in both health and diabetic retinopathy conditions while also interacting and organizing physiologically with iPSC-derived retinal pericytes. Our studies establish functional human iRECs and microphysiological iBRB models that facilitate mechanistic studies aimed at identifying therapeutic targets and promoting the revascularization of injured retinas, thereby supporting treatment advancement.

## Introduction

The development of human retinal vasculature begins at the fourth month of gestation and matures by the fifth month after birth^1^. Two major vascular systems, choroidal vasculature and inner retinal vasculature, are responsible for supporting outer and inner retinal barriers, respectively^2^. Retinal tissue has the highest energy and oxygen usage in the body due to the retina’s intense and continuous neuronal activity^3–5^. This demand leads to a crucial reliance on the inner blood-retina barrier (iBRB) to maintain ocular homeostasis^6,7^.

The iBRB is a specialized anatomical unit composing the vasculature located within the internal retinal layers of the central nervous system^7,8^. Retinal endothelial cells (RECs) in the iBRB are continuous endothelial cells (ECs) that form tight junctions to regulate the diffusion of small molecules, such as ions and water, across their cell-cell interface^9^. To develop the REC barrier, angiogenic and angiocrine signaling factors direct precursor RECs across the retina’s superficial plexus and into the deep retina tissue through guided angiogenesis^10–12^. These precursor RECs subsequently differentiate and mature under the guidance of Wnt/β-catenin, primarily driven by Norrin/Frizzled4 (Fz4), Sonic Hedgehog (Shh), and Notch signaling. Continued molecular signaling events and cellular interactions further drive the formation of specialized RECs^13–27^. Importantly, the disruption of the Norrin/Fz4 pathway abrogates retinal vascular development and iBRB functionality, suggesting its significance to REC development and retinal health^28–31^.

iBRB dysfunction and vascular leakage have been identified as a hallmark of numerous retinal microvascular diseases^7,32,33^, including diabetic retinopathy (DR), the leading cause of blindness and a common pathology found in patients with diabetes mellitus^33–35^. The breakdown of the iBRB occurs due to prolonged hyperglycemia exposure, leading to increased permeability of the endothelial barrier and reduced oxygen delivery to the retina, causing ischemia. As DR progresses, it can eventually lead to bleeding, retinal detachment, and irreversible blindness^36–38^. Despite DR’s significant health impact, our understanding of iBRB breakdown causality remains limited, severely constraining treatment options.

Human induced pluripotent stem cells (hiPSCs) have inexhaustible self-renewal and proliferation ability. They can differentiate into all three germ layers where they can be further directed into specialized cell types. So far, ophthalmic diseases have accounted for a significant portion of hiPSC clinical trials, reaching up to 24.4%^39^. The first clinical trial with hiPSCs was conducted on a patient with age-related macular degeneration^40^, a retinal microvascular disease. In the study, retinal pigment epithelial cell sheets from autologous hiPSCs were transplanted into the patient. Although the outcome showed no improvement, the patient’s condition remained stable for one-year post-treatment, highlighting the potential of hiPSC-derived cells for treating the eye.

In this study, we harnessed Wnt/β-catenin signaling, including the Norrin/Fz4 axis, to derive RECs from hiPSCs (iRECs) that can generate a continuous endothelial barrier with a characteristic retinal phenotype and genotype, as well as iBRB functionality. To investigate retinal microvascular diseases, we examined the response of the iRECs to an elevated glucose concentration and oxygen deprivation, representative of DR, and confirmed their recapitulation of clinical presentations. We established the therapeutic potential of the iRECs for the ischemic eye using an oxygen-induced retinopathy (OIR) mouse model. Finally, we synthesized *in vitro* human iBRB models, generating a perfusable vascular plexus that can be further supported by retinal pericytes. The robust and therapeutically active iRECs can model the iBRB in health and disease, providing a reliable source to elucidate pathophysiological mechanisms that improve treatments and enable precision medicine.

## Results

### Derivation of retinal endothelial cells from human induced pluripotent stem cells via the Norrin/Fz4 axis

To obtain RECs from hiPSCs, we first examined factors reported to be crucial in retinal vascular development. Norrin is identified as an essential and specific ligand that binds to Fz4 receptors on ECs to regulate retinal vascular development^28,41^. As a result of this pertinent role, vessel leakage was observed in Norrin-knock-out mouse retina^18^. It has also been reported that pericytes secrete vitronectin to induce and maintain REC barrier function during retinal vascular development^42^. Similarly, RepSox has been shown to enhance barrier function in human retinal microvascular ECs by inhibiting TGFβ/ALK5 and activating the REC-specific Notch and Wnt signaling pathways^43^. Based on this information, we incorporated these essential elements during EC differentiation and investigated their ability to generate retinal-specific ECs.

Mono-allelic mEGFP-tagged TJP1 WTC (WTC-TJP1) hiPSCs were differentiated by mesodermal induction using Essential 6 media supplemented with glycogen synthase kinase 3 (GSK3) inhibitor CHIR99021 for 48 hours^44–46^. Given the role of ETV2 in endothelial development,^47^ we used a previously established approach to differentiate the mesodermal cells into ECs by inserting chemically modified ETV2 mRNA using cell electroporation^48^ (**Fig 1A**). Following electroporation, cells were cultured in media supplemented with VEGF and bFGF for the EC fate^44–46,48^ as well as with Norrin and vitronectin for the pre-iREC specification. To produce non-tissue specific ECs as a control, referred to as iECs, electroporated cells were cultured in media supplemented with only VEGF and bFGF^44–46^ (**Fig S1A**).

**Figure 1.**
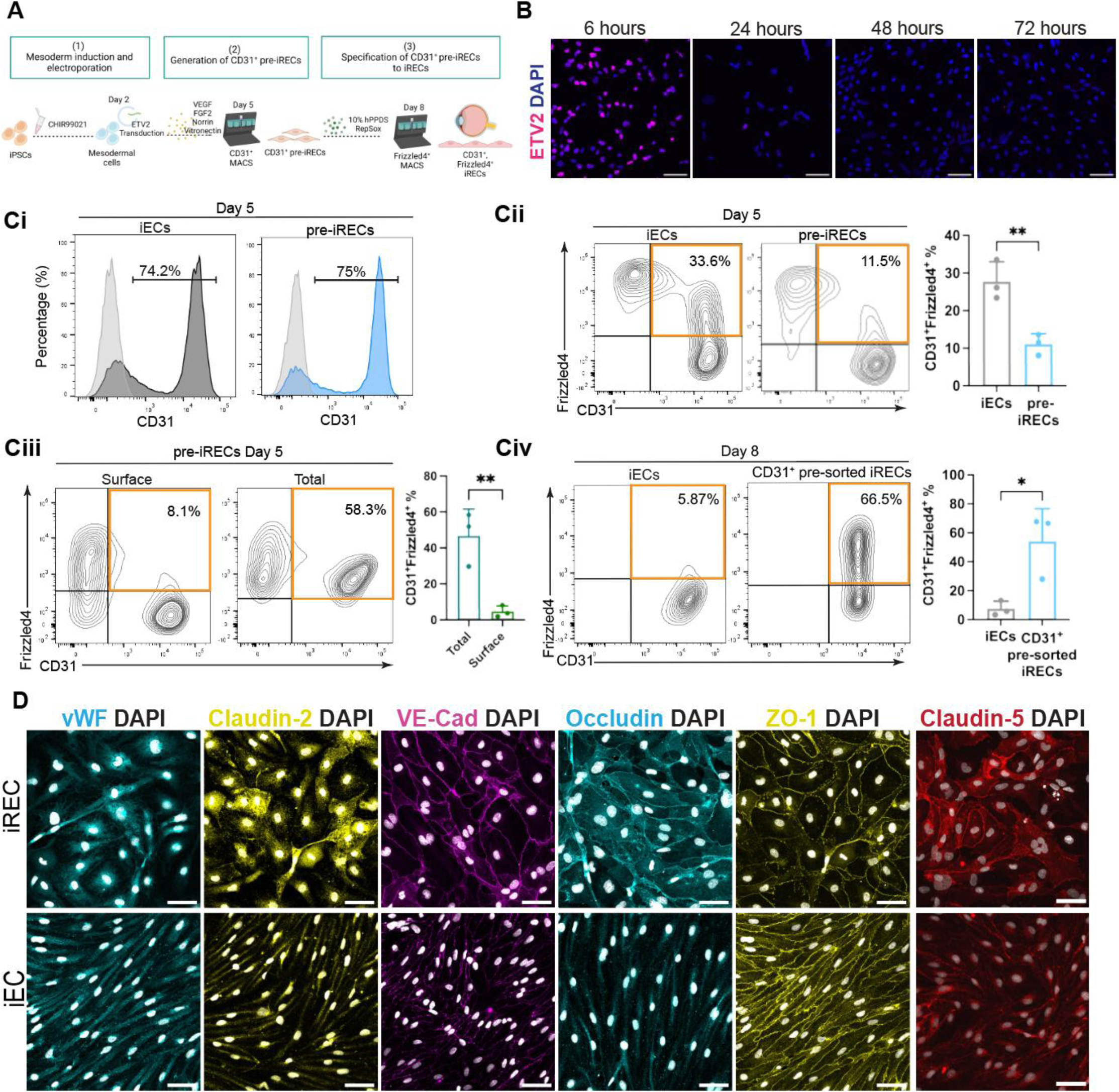
Derivation of retinal endothelial cells from human induced pluripotent stem cells via the Norrin/Fz4 axis. (A) Schematic illustration of iREC differentiation. (B) Time course immunofluorescence staining for ETV2 after electroporation (ETV2 in magenta, nuclei in blue). (C) i. Representative flow cytometry analysis of day 5 CD31 expression on iECs and pre-iRECs. ii. Representative flow cytometry analysis of CD31 and Frizzled4 expression on iECs and pre-iRECs on day 5 (*left*) and corresponding quantification (*right*). iii. Representative flow cytometry analysis of CD31 and Frizzled4 total and surface expression on day 5 pre-iRECs (*left*) and corresponding quantification (*right*). iv. Representative flow cytometry analysis of CD31 and Frizzled4 expression on iECs and CD31^+^ pre-sorted iRECs on day 8 (*left*) and corresponding quantification (*right*). (D) Immunofluorescence images of double sorted iRECs for tight junction protein and endothelial cell markers: vWF and Occludin (Cyan), Claudin-2 and ZO-1 (yellow), VE-Cadherin (magenta), Claudin-5 (red), and all nuclei in gray. Scale bars: 50 µm (B and D).

The robust but transient activation of ETV2 after electroporation (**Fig 1B** and **Fig S1B**) resulted in efficient endothelial fate. On day five, approximately 74% of hiPSCs expressed CD31 in both treatment groups (**Fig 1Ci**). In the pre-iREC group, the expression of β-catenin and cell junction marker ZO-1 emerged in parallel with the decrease of ETV2 expression (**Fig S1C**).

We next examined the expression of Fz4 as mouse, rat, and human studies have shown its specificity in RECs^43,49,50^. However, despite having high CD31 expression in both groups, we noticed that the addition of Norrin in the culture media resulted in a decreased CD31^+^Fz4^+^ population in the pre-iRECs on day five (**Fig 1Cii**). As it has been reported that Fz4 undergoes endocytosis following ligand-receptor binding^51^, we compared surface and total Fz4 expression via flow cytometry. Intriguingly, in around 50% of the pre-iRECs, Fz4 receptor was endocytosed (**Fig 1Ciii**), which suggests activation of the Norrin/Fz4 signaling pathway^52^.

After confirming the robustness and efficiency of the EC differentiation, we sorted CD31^+^ cells on day five. CD31^+^ pre-iRECs were then cultured in media with 10% human platelet-poor derived serum and RepSox for an additional three days (**Fig 1A**). In contrast, control CD31^+^ iECs were cultured in media with VEGF and TGF-β inhibitor, as previously reported^44–46^. Flow cytometry analysis was performed on day eight to determine if Fz4 was still detectable on both EC types. We observed that ∼66% of the CD31^+^ pre-sorted iRECs expressed Fz4 while a minimal amount of the iECs had the Fz4 receptor (**Fig 1Civ**). We further performed qRT-PCR and functional analyses to compare day eight iECs and CD31^+^ pre-sorted iRECs. Although CD31^+^ pre-sorted iRECs expressed higher levels of *TJP1* (ZO-1) and *OCLN* (occludin) (**Fig S1D)**, cell transendothelial electrical resistance (TEER) was comparable between these two groups (**Fig S1E)**. As only 66% of CD31^+^ pre-sorted iRECs expressed Fz4, we were concerned that a heterogeneous population of iRECs and non-specific iECs was present, impacting the results of the functional analyses and thus limiting the utility of this differentiation pathway.

To further enrich the REC phenotype and generate a pure REC population, we performed a second round of sorting on day eight. CD31^+^ pre-sorted iRECs were sorted for Fz4 to obtain CD31^+^Fz4^+^ double positive ECs (**Fig 1A**, referred to as iRECs). Immunofluorescence analysis showed that EC markers CD31, Ulex europaeus agglutinin I (UEA1), VE-Cadherin, Claudin-5, and vWF were detectable in both iECs and iRECs (**Fig 1D** and **Fig S2A**). Junctional proteins commonly found in RECs, including ZO-1, Occludin, and Claudin-2^7,53,54^ were highly expressed at the cell-cell junction in iRECs, while Occludin and Claudin-2 were not present to any significant degree in iECs (**Fig 1D**). We also examined the expression of EC and cell-cell junction markers in primary human RECs (HRECs, **Fig S2A** and **S2B**). HRECs express EC markers such as CD31, UEA1, vWF, and VE-Cadherin but not Claudin-5. Cell-cell junction markers, including Occludin, Claudin-2, and ZO-1 are hardly detectable in HRECs. This may be due to the tendency of primary cells to lose their phenotypic characteristics when removed from their original microenvironment. We were further able to demonstrate that iRECs derived from the C1-2 hiPSC line^55^ also possessed essential canonical EC and REC specific markers after undergoing the same iREC differentiation protocol (**Fig S2C**), emphasizing its robustness and utility.

Taken together, these results confirm that we can generate RECs from hiPSCs by activating the Wnt/β-catenin molecular signaling pathway via the addition of Norrin and vitronectin. The maturation of iRECs is further enhanced by incorporating the TGFβ/ALK5 inhibitor and Notch and Wnt signaling activator, RepSox, into the differentiation scheme.

### iRECs exhibit functionality and network properties typical of both healthy and diabetic retinopathy iBRB

Next, we examined if iRECs possessed functional characteristics fundamental to RECs found *in vivo*, including high barrier functionality, transcellular transport, and the ability to form vascular networks. We observed higher TEER in the iRECs relative to the iECs (**Fig 2A**), displaying enhanced barrier properties. As a consequence of the iBRB’s highly selective permeability, RECs utilize transcellular transport as the primary mechanism for delivering and removing molecules from the brain and retinal parenchyma^7,56^, making this a key functionality that RECs must possess. The glucose transporter 1 (GLUT1) receptor plays a crucial role in the retina. The GLUT1 transporter is the main facilitator of glucose transport, which may have significant implications for DR^57,58^. When the iRECs were treated with the highly selective GLUT1 inhibitor, Bay-876, glucose uptake was significantly reduced (**Fig 2B**). Taken together, these results demonstrate that the iRECs possess functional barrier and transporter activity.

**Figure 2.**
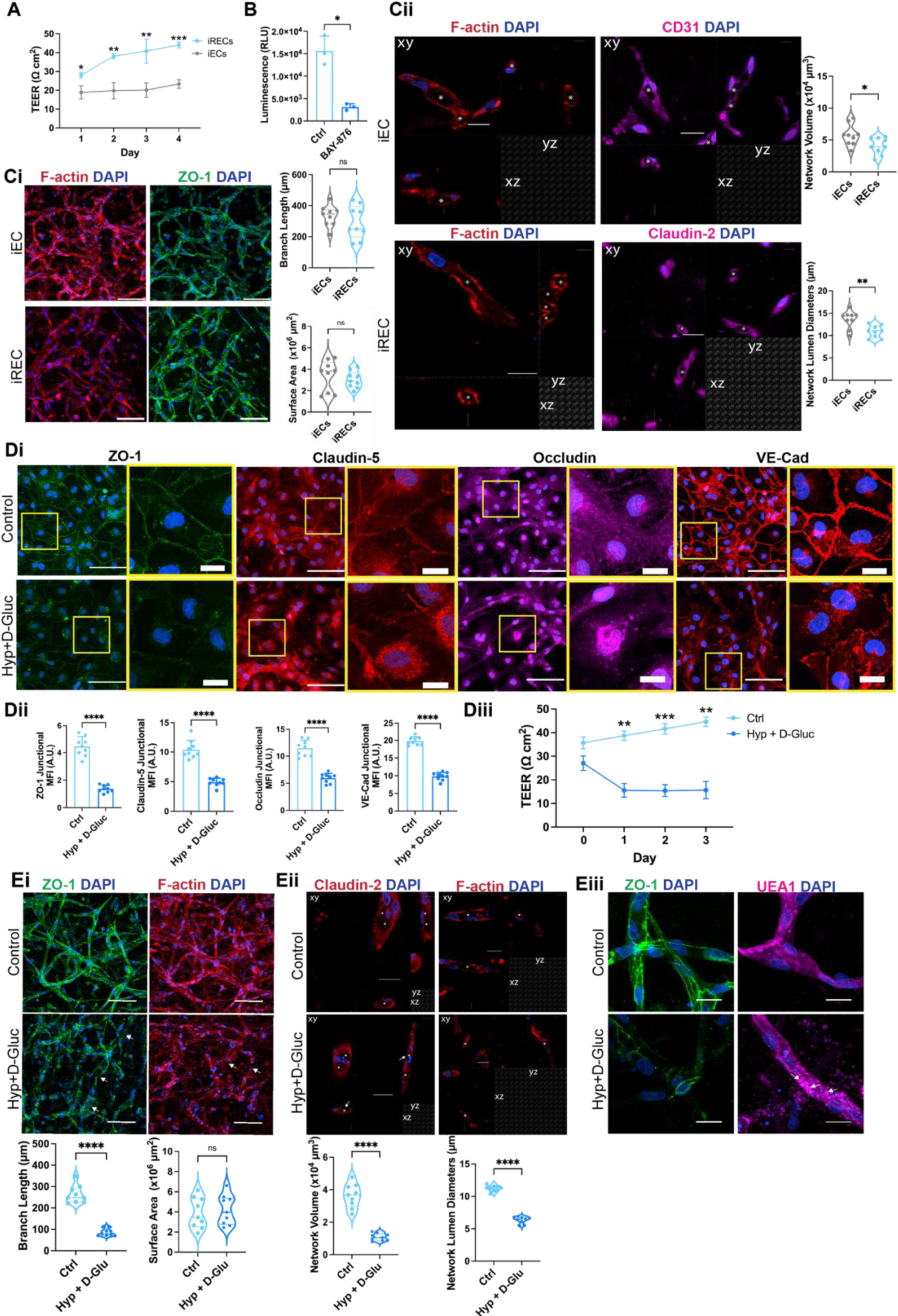
iRECs exhibit functionality and network properties typical of both healthy and diabetic retinopathy iBRB. (A) TEER measurements of iECs and iRECs. (B) GLUT1 uptake assay for iRECs treated with or without the GLUT1 inhibitor, Bay-876. (C) i. Representative immunofluorescence images of 3D iEC and iREC vascular networks (*left*) and quantification of their branch lengths and surface areas (*right*). F-actin (red), ZO-1 (green), and nuclei (blue). ii. Cross-sectional views of 3D iECs and iREC vascular networks depicting their lumina (asterisks) and expression of F-actin (red) and CD31 or Claudin-2 (magenta; *left*). Quantification of vascular network volumes and lumen diameters (*right*). (D) i. Representative immunofluorescence images of junctional protein markers on iRECs in control or diabetic conditions (Hyp + D-Gluc). ZO-1 (green), Claudin-5 and VE-Cad (red), Occludin (magenta), and nuclei (blue). ii. Corresponding quantification of mean fluorescent intensity at cell-cell junctions. iii. TEER measurements of iRECs with or without diabetic treatment (E) i. Representative immunofluorescence images of 3D iREC vascular networks with or without diabetic treatment (*top*) and quantification of their branch lengths and surface areas (*bottom*). F-actin (red), ZO-1 (green), and nuclei (blue). Arrows depict examples of disrupted vasculature. ii. Cross-sectional views of 3D iREC vascular networks with or without diabetic treatment, illustrating altered morphology in the diabetic condition (*top*), and quantification of network volumes and lumen diameters (*bottom*). F-actin and Claudin-2 (red) and nuclei (blue). Asterisks denote lumens and arrowheads highlight disconnected lumens. iii. Representative immunofluorescence images of 3D iREC vascular networks expressing ZO-1 and UEA1 with or without diabetic treatment. Arrowheads highlight granular objects present within the UEA1 stain. ZO-1 (green), UEA1 (magenta), and nuclei (blue). Scale bars: 100 µm (Ci, Di originals, and Ei) and 20 µm (Cii, Di insets, Eii, and Eiii).

The capability of ECs to form vascular networks that contain lumens is an essential aspect of their physiological properties and is thus a characteristic requirement for iRECs. In 3D collagen type-I hydrogels, iRECs robustly formed capillary-like structures. While their vascular branch length and surface area were consistent with iECs, they had smaller overall network volumes and narrower lumen diameters relative to iECs (**Fig 2C**), consistent with values measured in humans^59,60^. The iREC capillary-like networks also expressed key canonical endothelial cell and junctional protein markers. Specifically, the iREC networks contain heightened membrane-bound Claudin-2 and Claudin-5 cell-cell contact expression relative to iEC networks (**Fig S3A**), while CD31, ZO-1, VE-Cadherin, β-catenin, UEA1, and F-actin expression patterns did not appear to differ between iREC and iEC networks (**Fig S3B**), supporting an enhanced iREC phenotype and endothelial barrier stability. Cross-sectional imaging highlights the presence of physiological lumina within the iREC networks that are adhered together at cell contacts via the Claudin-2 junctional protein (**Fig 2Cii**). Alternatively, cell contacts in iECs networks could be observed via CD31 rather than Claudin-2, as their limited Claudin-2 cell-cell contact expression confounds lumen and network delineation and identification (**Fig 2Cii and S3A**). Taken together, this data confirms the accuracy, specificity, and efficacy of the iREC differentiation and the robustness of our models, critical features that will enable accurate and translatable REC-focused research.

In proliferative DR, hypoxia and hyperglycemia have been implicated as causative factors, inducing microvascular degeneration that coincides with decreased EC barrier function and disrupted cell-cell junctions^7,33,34,61–65^. To recapitulate microvascular changes in DR, we exposed iRECs to a high concentration of D-glucose (30 mM) in hypoxia (1% O_2_). After 24 hours of incubation, tight junction proteins ZO-1, Claudin-5, and Occludin as well as the adherens junction protein VE-Cadherin, displayed decreased localization at cell-cell contacts (**Fig 2Di-ii**). Interestingly, the mean fluorescent intensity of Claudin-2 at the junctions did not change, suggesting that Claudin-2 does not appear to be impacted in DR (**Fig S4A**). To confirm that diabetic conditions did not induce unexpected phenotypic changes in iRECs, both conditions were stained for the EC biomarkers CD31 and UEA1, where they displayed consistent expression between culture conditions (**Fig S4A**). RT-qPCR analysis revealed no differences in the expression levels of *CLDN5* (Claudin-5), *OCLN* (Occludin), and *TJP1* (ZO-1) (**Fig S4B**). This observation further emphasizes that the initial response to diabetic conditions involves protein re-localization rather than impacting expression. Finally, TEER was examined after exposing iRECs to DR conditions for 72 hours to evaluate its influence on barrier properties. We observed that the diabetic-treated iRECs contained significantly lower TEER levels compared to the control, demonstrating reduced iREC barrier functionality, consistent with the clinical presentation of DR (**Fig 2Diii**). Next, we investigated the responses of iREC vascular networks to DR conditions. We found that while the retinal vascular networks exposed to DR conditions did not display a statistically significant difference in vascular surface area, they had much smaller branch lengths, network volumes, and lumen diameters (**Fig 2Ei-ii**). The consistency in surface area across conditions may be due to the DR phenotype compensating for area with an increase in smaller, pathological vasculature. ZO-1 and F-actin stains display disconnected networks and cross-sections depict discontinued and aberrant lumina within the diabetic treated condition (**Fig 2Ei-ii**). Furthermore, after DR treatment, junctional protein expression, including ZO-1, CD31, VE-Cadherin, and Claudin-5 decreased at cell-cell contacts, but Claudin-2 remained unchanged (**Fig 2Eiii and S4C**), suggesting a loss in barrier functionality within the iREC networks. Interestingly, under DR conditions, UEA1 granular objects appeared in the 3D vascular networks (**Fig 2Eiii**), suggesting glycocalyx degradation that has been previously reported in diabetic patients^66,67^.

In summary, we demonstrate that iRECs possess the functional characteristics of RECs with responsiveness to changes in glucose and oxygen. As a result, they contain the properties to serve as a reliable and robust source for cell-based therapeutics and *in vitro* disease modeling platforms.

### Transcriptomic analysis of the iRECs reveals their distinct recapitulation of the REC genotype

To elucidate the transcriptional profile of iRECs, we performed a bulk RNA sequencing (RNA-Seq) analysis three days after Fz4 sorting (i.e. CD31^+^ Fz4^+^). We used iECs (i.e. CD31^+^) to assess the tissue-specific traits of iRECs. Principle component analysis (PCA) of differentially expressed genes showed that the iRECs and iECs cluster distinctly apart from each other as there was only transcriptional similarity across biological replicates within a phenotypic condition (**Fig 3A**). This result is further confirmed by correlation analysis (**Fig S5A)** and a heat map of hierarchical clustering analysis of differentially expressed genes based on enriched Gene Ontology (GO) categories (**Fig 3B**), indicating genetic differences between iRECs and iECs.

**Figure 3.**
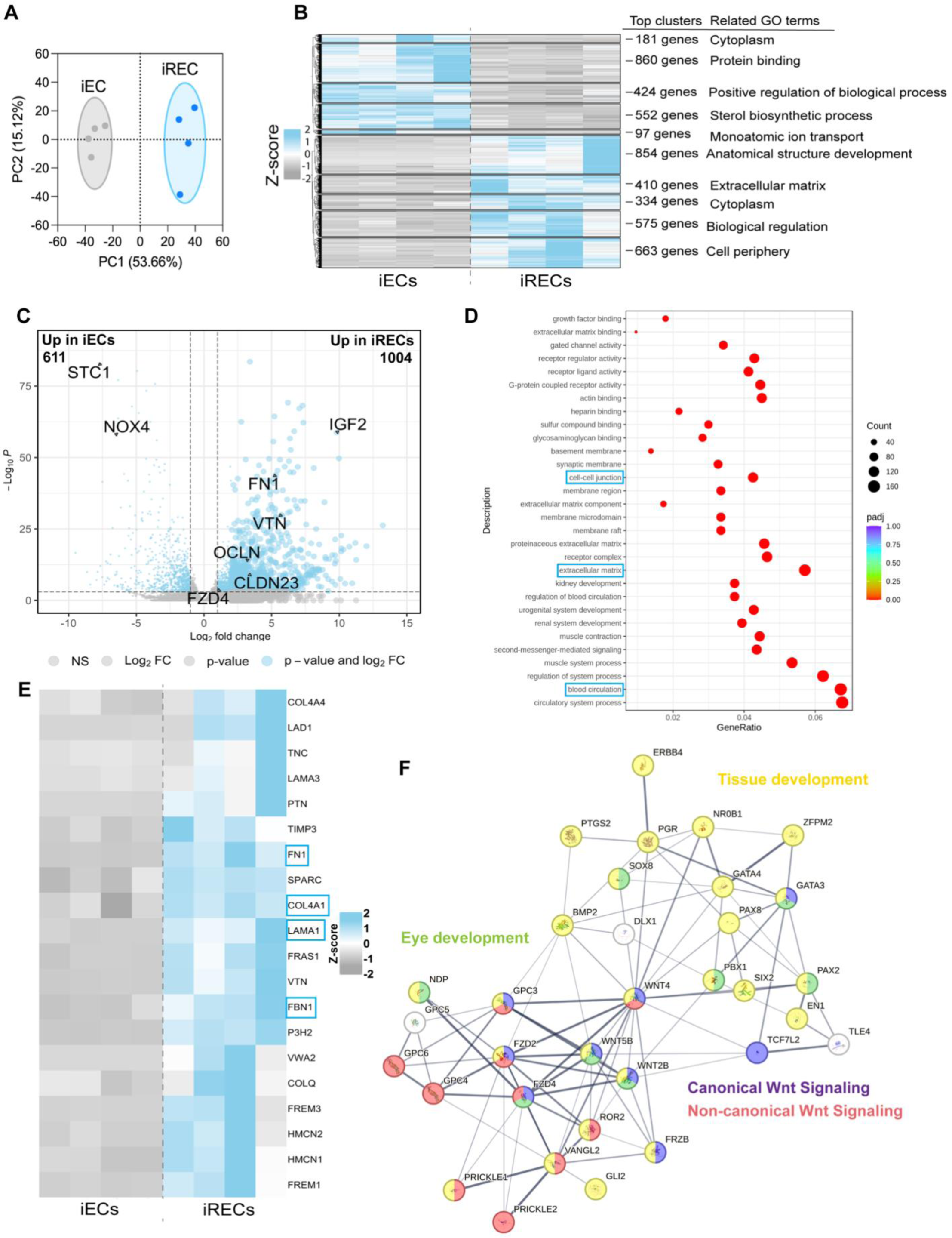
Transcriptomic analysis of the iRECs reveals their distinct recapitulation of the REC genotype. RNA-Seq data analyses from four biological replicates (independent differentiation) of iECs and iRECs, including (A) Principal component analysis, (B) Heat map analysis of differential gene expression (FC>2), (C) Volcano plots of differentially expressed genes (FC>2), (D) GO analysis (total enrichment), (E) Heatmap analysis of basement membrane-related genes (FC>2), and (F) Network analysis of Wnt signaling pathways involved in the generation of iRECs.

Focusing on known gene differences, we confirmed that iECs and iRECs shared 11238 genes while 1476 genes were distinctly expressed in iRECs (**Fig S5B**). Compared to iECs, 1004 differentially expressed genes were significantly upregulated in iRECs, including those closely associated with REC function, such as *FZD4* (1.15 Log2 FC), *IGF2* (9.8 Log2 FC), *CLDN10* (6.54 Log2 FC), *CLDN23* (3.32 Log2 FC), *VTN* (5.71 Log2 FC) and *FN1* (5.29 Log2 FC; **Fig 3C**). To further understand the transcriptional difference between iECs and iRECs, we performed a detailed GO enrichment analysis (**Fig 3D**). Significant enrichment of genes associated with extracellular matrix, blood circulation, and cell-cell junction was found in iRECs compared with iECs, suggesting that the processes related to cell remodeling, cellular adhesion, and endothelial cell function, are prominently activated in iRECs compared with iECs.

We next analyzed the transcriptomic factors contributing to the enhancement of iREC phenotypes. We generated heat maps to examine differential gene expression associated with the basement membrane and cell-cell junctions (**Fig 3E and Fig S6A**). Notably, basement membrane components *COL4A1* (Collagen IV), *FN1* (Fibronectin), *LAMA1* (Laminin-1), and *FBN1* (Fibrillin-1) are upregulated in iRECs. Within cell-cell junction proteins, *OCLN* (Occludin), *CDH6* (Cadherin-6), and *CLDN23* (Claudin-23) are highly expressed in iRECs compared with iECs. These genes are involved in REC development and barrier formation^34,54,68–70^, supporting the genotypic characteristics and phenotypic integrity of the iRECs.

Interested in the potential pathways involved in the generation of the iRECs, we investigated the Wnt pathway as it is implicated as a significant contributor to the development of RECs and iBRB formation^70,71^. Dysregulation of Wnt signaling has been associated with retinal diseases, such as familial exudative vitreoretinopathy and Norrie disease^72^. As a result, Wnt activation is fundamental to RECs and retinal health and thus must be possessed by the iRECs. To this end, we generated heat maps to highlight differential gene expression associated with the Wnt signaling pathways (**Fig S6B**). The major Wnt signaling pathway components *FZD4* (Fz4), *LRP5* (low-density lipoprotein receptor-replated protein 5), and *APCDD1* (APC down-regulated 1) were upregulated in iRECs compared with iECs. Network analysis reveals the involvement and the increased expression of both canonical and non-canonical Wnt signaling pathways, including Wnt4, Wnt5b, and Wnt2b (Wnt13) in iRECs compared to iECs (**Fig 3F**). Notably, *FZD4, LRP5* and *NDP* (Norrin) have been implicated in REC differentiation and iBRB formation^20^. *VANGL2* (VANGL planar cell polarity protein 2) and *TCF7L2* (Transcription factor 7 like 2) control cell fate decisions during REC development^73,74^. Wnt2b (Wnt13) interactions with both Wnt4 and Wnt5b are involved in vascular formation and vasculogenesis^72,75^. Although the direct relation of Wnt2b (Wnt13) with retinal cell differentiation is not well-studied, its involvement in the canonical Wnt pathway in modulating peripheral fate of the chick eye suggests it could indirectly influence EC development and iBRB formation in the retina^76^, supporting its activation within the iRECs and thus their robustness.

Given that both brain ECs and RECs originate from the central nervous system, it is reasonable to speculate that they may share similar genotypes due to their analogous function. To elucidate this assumption, we conducted a comparative analysis of published RNA-Seq data of iPSC-derived brain ECs^77^, compared to our iRECs and iECs. Both correlation and PCA analysis showed that all cell types cluster distinctly from each other and are minimally correlated (**Fig S7A-B**). The number of distinctly expressed genes among iRECs, iECs, and iPSC-derived brain ECs was also investigated (**Fig S7C**), revealing shared EC traits and tissue-specific differences between the groups.

In conclusion, these data highlight the distinct transcriptome of iRECs, the signaling pivotal to the development of RECs, and the iREC’s recapitulation of fundamental and developmental REC markers. This displays the biological fidelity of the iRECs, the developmental insight they can provide, and the utility they possess.

### iRECs revascularize the ischemic eye

To investigate the therapeutic potential of iRECs in treating DR, we used the OIR mouse model, the most widely used model for studying DR angiogenesis/neovascularization (NV) and the discovery and testing of therapeutics^78,79^. We sought to examine whether implantation of iRECs could rapidly revascularize the ischemic eye and thus regenerate and repair the tissue. To do this, we intravitreally injected iRECs into one eye of the OIR mice and PBS into the contralateral eye as the vehicle control on postnatal day 12 (P12). We investigated vaso-obliteration (VO) and NV area on P17, as this is when they are at their peak within the OIR model^80,81^. Both VO and NV were significantly reduced in the treated eye compared to the contralateral eye injected with PBS (**Fig 4A-B**), demonstrating the reparative and therapeutic impact of the iRECs.

**Figure 4.**
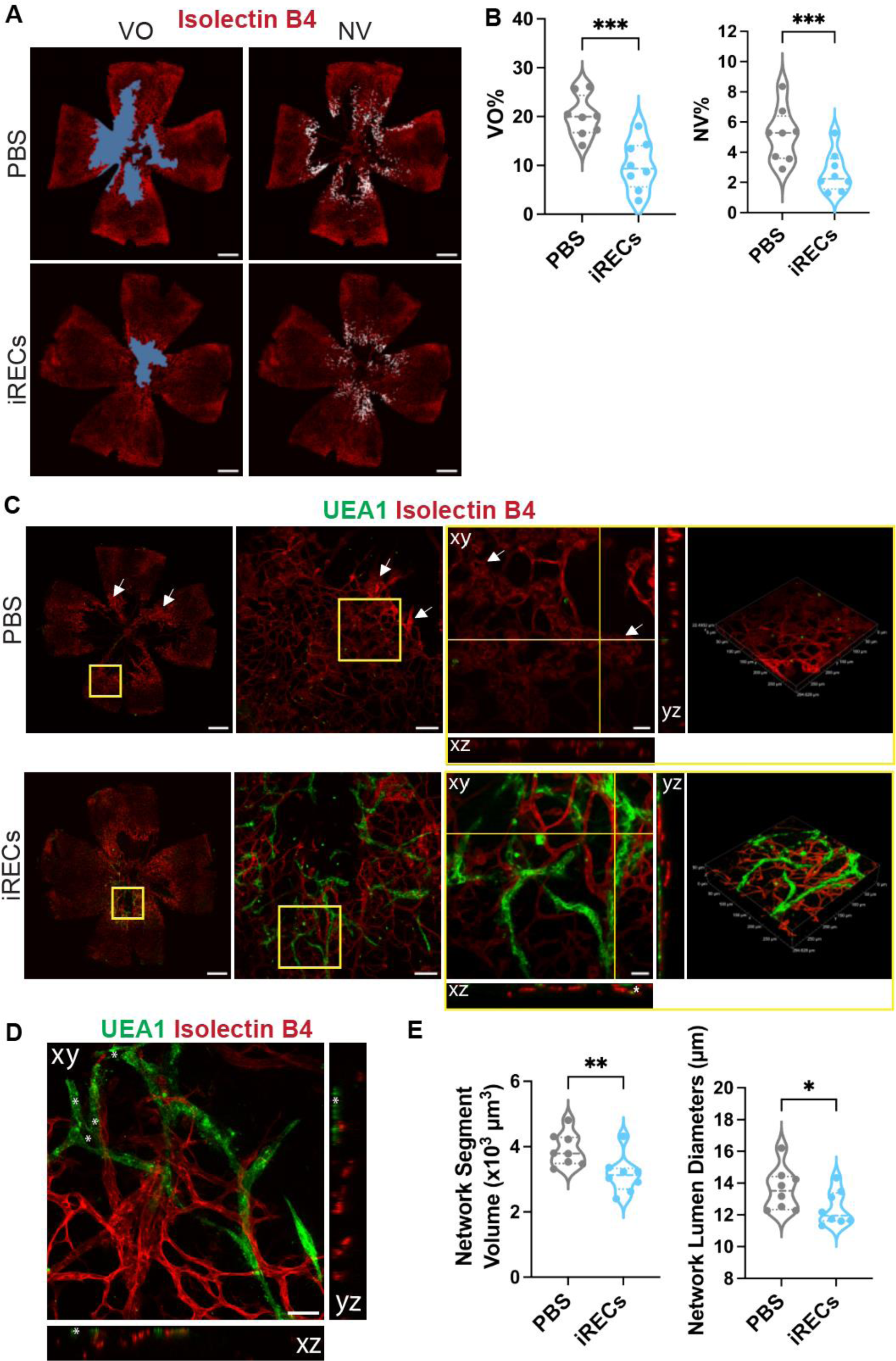
iRECs revascularize the ischemic eye. (A) Representative immunofluorescence images of the OIR mouse retinal tissue at postnatal day 17 (P17), five days after intravitreal injection of PBS or iRECs. Isolectin B4 (magenta), vaso-obliteration VO area (blue), and pathological neovascularization (NV) area (white). Scale bars: 300 µm. (B) Quantification of VO and pathological NV at P17. (C) Representative immunofluorescence images of whole mount retina, cross-sections of the OIR mouse retinal tissue, and 3D reconstructions of mouse vascular regions at P17 for PBS (*top*) and iREC (*bottom*) intravitreal injections. Mouse Isolectin B4 (red) and human UEA1 (green). Arrows depict pathological neovascular tufts and yellow crosshairs demonstrate location of xz and yz planes. Scale bars: 500 µm, 100 µm, and 25 µm, (left to right). (D) An immunofluorescence cross-sectional image of lumenized iREC vascular networks integrating with OIR mouse vasculature tissue on P17. Mouse Isolectin B4 (red) and human UEA1 (green). Asterisks denote lumens of human/mouse hybrid vasculature. Scale bar: 50 µm. (E) Quantification of mouse vascular network segment volume and lumen diameter across PBS and iREC injected groups. All quantifications are n = 8.

After establishing the ability of the iRECs to restore vascularization and rescue the ischemic eye, our next step was to identify if the iRECs actively participated in this process.

To do this, we monitored cell integration with the host on P17 just before healthy vascular structure begins to spontaneously form and mature in the OIR retina^82,83^. We observed that iRECs integrated with the host vasculature. Specifically, we noted that iRECs formed complex vascular networks in the hypoxic, avascular regions of the mouse retina, while the PBS control retina contained pathological vasculature with significant VO and neovascular tufts (**Fig 4C**). These networks were lumenized, consisting of human-only and human-mouse hybrid networks, indicating vasculogenesis of the iRECs and their integration with the host vasculature (**Fig 4C-D**). Exploring the impact of these cellular interactions, we quantified the mouse vascular networks exposed to treatment and control. While the branch length and surface area were not different (**Fig S8A**), we found that the mouse vascular networks within the iREC group had smaller segment volumes and lumen diameters than their control counterparts (**Fig 4E**). The smaller networks in the iREC condition more closely align with healthy murine retinal capillary values^84,85^, suggesting a decrease in NV and pathological alterations relative to the PBS group. Interestingly, the PBS group appears to have some iRECs present within the mouse retinal tissue, including the interior of an artery where they may be migrating (**Fig 4C and S8B**). This demonstrates that the injected iRECs are able to transport to and potentially treat the contralateral eye. Corroborating these findings, a gene therapy study has reported the bilateral transfer of the substances between eyes, with visual improvement also observed in the contralateral, non-treated eye^86^. Therefore, we speculate that the consistency in branch length and surface area across groups may be a consequence of iREC infiltration within the control. Overall, the vasculogenic and angiogenic capabilities of the iRECs lead to the restoration of the ischemic eye, suggesting that they could be used as a strategy to repair the damaged retina.

### iPSC-derived iBRB-on-a-chip

Microphysiological systems (MPS), or organs-on-chips, are downsized, functional *in vitro* models that faithfully recapitulate human organs, enabling the study of tissue function in health and disease^87–108^. We thus sought to establish the capacity of the iRECs to generate a functional iBRB-MPS. To develop the MPS, we utilized a standard PDMS-based three-cell channel microfluidic device^109^. iRECs encapsulated within a collagen type-I hydrogel were delivered into the cell culture channels in the fluidic device and cultured for three days to induce the formation of iREC vascular networks (**Fig. 5A**). In the MPS, iRECs self-assembled into 3D networks that contained canonical membrane-bound adherens and tight junction proteins throughout the networks and at cell-cell contacts (**Fig. 5B-C, S9A**), characteristic of the functional iBRB. Cross-sectional images show the presence of lumina that recapitulate iBRB morphology and are adhered at cell contacts via the Claudin-2 junctional protein (**Fig. 5Di**). Further corroborating this, we flowed 70 kDa rhodamine dextran through the iREC networks, demonstrating their perfusability and the presence of continuously connected open luminal networks (**Fig. 5Dii, S9B**).

**Figure 5.**
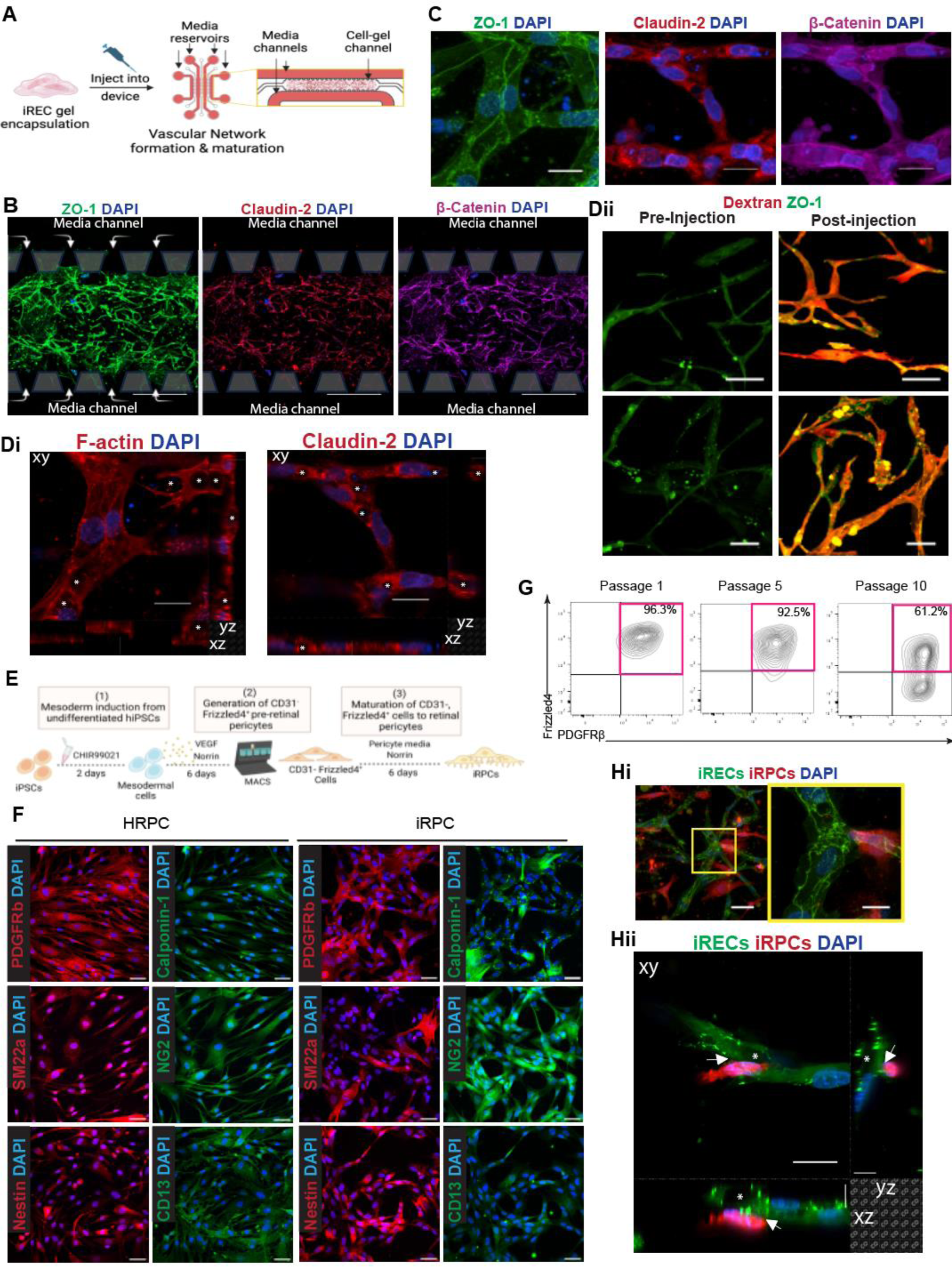
iPSC-derived iBRB-on-a-chip. (A) Schematic depicting the development of the iBRB-on-a-chip with iRECs. (B) Representative immunofluorescence images of 3D iREC vascular networks within the MPS. ZO-1 (green), Claudin-2 (red), β-catenin (magenta), and nuclei (blue). Images are overlaid with markings of the channel posts and the location of the media channels. Arrows depict the flow of media into the cell-gel channels. (C) Representative immunofluorescence images of 3D iREC vascular networks within the MPS expressing tight and adherens junction proteins at cell-cell contacts. ZO-1 (green), Claudin-2 (red), β-catenin (magenta), and nuclei (blue). (D) i. Cross-sectional views of 3D iREC vascular networks within the MPS showing lumenized vessels (asterisks) and expression of the tight junctional protein, Claudin-2, at cell-cell contacts. F-actin and Claudin-2 (red) and nuclei in blue. ii. Representative immunofluorescence images of the 3D iREC networks before (*left*) and after (*right*) 70 kDa rhodamine dextran injection. ZO-1 (green) and rhodamine dextran (red). (E) Schematic illustration of iRPC differentiation. (F) Immunofluorescence images of pericyte markers in HRPCs (left) and iRPCs (right). PDGFRβ, SM22α, and Nestin (red), calponin-1, NG2, and CD13 (green), and nuclei (blue). (G) Flow cytometry analysis of Fz4 and PDGFRβ across iRPCs at passages one, five, and ten. (H) Representative immunofluorescence images of the iREC-iRPC co-cultured iBRB-on-chip displaying i. perivascular iRPCs (red) surrounding and encapsulating iREC networks (green) and ii. an iRPC (red) interacting with and supporting an iREC network lumen (green). Asterisks denote lumens and arrows highlight iRPC-iREC interactions. Scale bars: 500 µm (B), 100 µm (Dii top panel), 50 µm (Dii bottom panel, F, and Hi left), 20 µm (C, Di, Hi right, and Hii xy plane), and 10 µm (Hii xz and yz planes).

Pericytes are critical for the development and maintenance of iBRB functionality and contribute to iBRB breakdown in disease^14,16,27,41,42,110–116^. We therefore sought to include pericytes in the iBRB MPS to increase the complexity and biological relevance of our model. To this end, we utilized an established pericyte differentiation protocol^44,117^ where pericytes are derived from iPSCs through a mesodermal lineage induction. As it has been suggested that Norrin/Fz4 signaling interactions are critical to iBRB development and retinal pericytes express Fz4^18–20,26,28,41,118,119^, we utilized Norrin supplementation during the differentiation to specifically develop iPSC-derived retinal pericytes (iRPCs; **Fig. 5E**). To characterize the iRPCs and validate their phenotype, immunofluorescent assays and flow cytometry analysis were performed. Immunofluorescence analysis demonstrated that the iRPCs properly expressed the essential PC markers PDGFRβ, SM22α, NG2, Calponin-1, CD13, and Nestin (**Fig. 5F**). Their expression is consistent with primary human RPCs (HRPCs), supporting the phenotypic integrity and biological accuracy of the iRPCs. The iRPC population also expressed the retinal PC marker, Fz4^41^, in conjunction with PDGFRβ, through passage 10, suggesting the retinal specificity of the PCs. Specifically, over 92% of the iRPC population co-expressed Fz4 and PDGFRβ after passage 5, with a substantial 61.2% of the population still expressing Fz4 and PDGFRβ at passage 10 (**Fig. 5G**). This validates the purity and longevity of the iRPCs and establishes the importance of the Norrin/Fz4 signaling axis to retinal pericyte development. After confirming the robustness of the iRPCs, we developed an iBRB MPS with both retinal-specific ECs and pericytes. We observed physiological iREC-iRPC interactions within the iBRB MPS (**Fig. 5Hi**), namely the formation of luminal iREC structures supported and encapsulated by iRPCs (**Fig 5Hii and S9C**).

Taken together, we show that our iPSC-derived iBRB model contains high biological fidelity, recapitulating physiological endothelial-pericyte organization and cellular interactions. This demonstrates that the iRECs and iRPCs can be utilized to generate a functional and perfusable iBRB MPS to study eye development and disease progression and thus advance therapeutics.

## Discussion

There is an increasing need to develop tissue-specific ECs for more precise disease modeling and effective tissue regeneration and repair. While primary human ECs can be utilized for these studies, they typically lose their *in vivo* characteristics when cultured *in vitro*^120–122^. This limits our ability to utilize these cells for understanding EC function during development or disease progression. RECs are unique as they possess a high barrier function that separates the central nervous system and circulatory system and regulates transport between them, ultimately supporting the functional eye and maintaining ocular homeostasis^6–8,123,124^. Retinal microvascular diseases impair iBRB function, thereby impacting vision^7,32,33^. As a result, developing biologically accurate and functionally active RECs is critical to understanding the iBRB in health and disease and advancing therapeutic discovery. Although iPSC-derived EC protocols have been developed^92,93,125–135^, including for iPSC-derived brain ECs^136–143^, methods to generate iPSC-derived RECs are not currently present. Transient activation of ETV2 has been shown to rejuvenate mature endothelial cells^144^ and induce endothelial development^47^. Therefore, we began by using a previously established approach of inserting chemically modified ETV2 mRNA into mesodermal cells to induce EC differentiation with high efficiency, resulting in shorter culture times and higher EC yield^48^. We then harnessed the Wnt/β-catenin signaling pathway, critical for REC development, during this EC differentiation to develop a novel approach for obtaining iRECs. Specifically, we identified that the Norrin/Fz4 axis and vitronectin signaling a part of the Wnt/β-catenin pathway are critical mediators of REC induction and are required for the development of RECs. Additionally, we established that the TGFβ/ALK5 inhibitor and Notch and Wnt signaling activator, Repsox, is required in the maturation of RECs. We thus elucidated that Norrin, vitronectin, and Repsox are signaling molecules necessary for the development of CD31^+^Fz4^+^ iRECs.

Next, we verified the phenotypic integrity and functionality of the iRECs, ensuring they contained the biological relevance to advance research and therapeutic discovery. The iRECs expressed key EC markers, including CD31, UEA1, VE-Cadherin, and vWF, as well as the retinal-specific EC markers Fz4 and Claudin-2^52,54^. They also contained high barrier and selective permeability properties, evidenced by their increased TEER relative to iECs, functional GLUT1 transporter activity, and expression of junctional proteins. Within a 3D hydrogel, the iRECs were able to self-assemble into vascular networks that recapitulated the morphology, junctional protein expression, and structure of the functional and mature iBRB^6,7^. Of particular importance, the iREC networks expressed membrane-bound Claudin-2, a unique marker found in RECs, but not brain ECs^54^, that has not been previously visualized at the protein level within human RECs. Taken together, this demonstrates that the Wnt/β-catenin signaling-derived iRECs possess physiological iBRB characteristics and functionality, displaying their ability to act as a robust and reliable source for iBRB and retinal microvascular disease research.

The leading cause of retinal microvascular disease and blindness, DR, is characterized by iBRB breakdown as a result of prolonged hyperglycemia exposure and ischemia^33–38,145^. To this end, we exposed the iRECs to hypoxia and high glucose to highlight the applicability of our iRECs in the pathological context, successfully recapitulating DR clinical presentations in both 2D and 3D. Barrier protein disruption was observed in both settings, with vascular regression and dysfunction found to be particularly significant in 3D culture. Specifically, ZO-1 and Claudin-5 displayed minimal junctional localization under diabetic conditions, consistent with previous work^146^. We also demonstrated a decrease in Occludin localization at cell-cell contacts under the DR condition. This contrasts with previous studies that exposed cells only to one diabetic feature, including iPSC-derived brain ECs to 1% O_2_^146^ or human primary RECs to 30 mM D-glucose and inflammatory cytokines^147^, which did not report changes in Occludin. In this study, we utilized both a high concentration of glucose and hypoxic conditions to effectively recapitulate DR, an approach that was not present in the other studies. Therefore, we speculate that synergistic interactions may account for these differences. Importantly, using these conditions, we were able to visualize UEA1 granular objects within 3D retinal vascular networks exposed to the DR conditions, pointing to glycocalyx degradation that has been reported clinically in diabetic patients^66,67^. Interestingly, we did not observe changes in Claudin-5, Occludin, and ZO-1 mRNA expression after a 48-hour exposure to DR conditions, despite observing reduced localization at cell-cell junctions in these conditions. This may suggest that the pathological changes associated with DR primarily affect protein stability or translation or post-translation modifications rather than transcription, warranting investigation in the future. Overall, we were able to robustly recapitulate the DR phenotype in 2D and 3D with our iRECs, exemplifying their ability to be utilized for *in vitro* disease modeling and to elucidate aberrant pathways and therapeutic targets.

To assess the biological fidelity of the iRECs at the genetic level and establish critical molecular signaling pathways associated with their development, we performed a transcriptomic analysis. We identified that iRECs contain a distinct genotype relative to iECs, including increased expression of the REC-specific *FZD4* and *LRP5* genes^52^, validating their specificity and thus the accuracy of the differentiation. The increased expression and involvement of Wnt signaling molecules in the iRECs relative to iECs further establishes the importance of Wnt signaling to REC development, identifying a fundamental requirement for iREC derivation. We next compared RNA-Seq data of iRECs with published iPSC-derived brain EC RNA-Seq data^77^. We found that while RECs and iPSC-brain ECs share a central nervous system origin, the two cell types have distinct gene expression profiles, indicating significant functional differences. These differences between the iPSC-derived brain ECs and iRECs should be examined more closely to identify the unique characteristics associated with each EC type. As it has been suggested that the transduction of ETV2 and related factors during brain EC differentiation assists in their endothelial fate and functionality^148^, comparing these derivatives to iRECs could provide more specific insights that elucidate retinal specific features. Overall, utilizing RNA-Seq, we confirmed our distinct recapitulation of RECs and uncovered signaling critical to their induction, validating their translatability and application to iBRB research and retinal vascular development studies.

To examine the therapeutic potential of the iRECs, we intravitreally injected them into an OIR mouse model for DR. The iRECs were able to integrate with host vasculature, especially within hypoxic, avascular regions, to reduce VO and NV and stimulate robust revascularization, markers of therapeutic success. In addition, the iRECs stimulated the formation of host vasculature volumes and lumen diameters that closely aligned with healthy murine retinal capillary values^84,85^, suggesting normalization of the vascular network, another indication of vascular network recovery. Thus, we demonstrate that the iRECs can rescue the ischemic eye, restoring normal retinal vascularization. Interestingly, we found that iRECs were able to migrate to the contralateral eye, undergoing a bilateral transfer previously reported in a gene therapy study^86^. This further displays the therapeutic capabilities of the iRECs, as they can transport to ischemic regions outside the injection site to help rescue the tissue. As this therapeutic impact takes place in immunocompetent C57B/6 mice, the iRECs possess clinical translatability and thus represent a promising treatment option.

Due to the significant biological mimicry of MPS, studies have shown their use in mechanistic studies and their potential to identify and test therapeutics, as well as accelerate protective countermeasure development^91,92,96,97,101,103,105,107,149,150^. To increase the complexity of our modeling, we developed a functional iBRB MPS. Utilizing self-assembly, we fabricated a perfusable retinal microvascular MPS that expresses canonical EC, REC, and junctional protein markers. Pericytes play an important role in iBRB formation and health in the retina^14,16,27,41,42,110–116^, yet their tissue-specific identity is not well understood. To integrate pericytes into the iBRB MPS, we first successfully generated retinal-specific pericytes from iPSCs that expressed canonical pericyte markers and the retinal-specific Fz4 via the incorporation of Norrin during cell differentiation. This identifies Norrin as a key modulator of RPC induction and a necessary signaling molecule for the development of iRPCs. Integrating the iBRB-on-a-chip with the iRPCs, we developed a functional iBRB MPS consisting of physiological endothelial-pericyte interactions and organization. Harnessing the mimicry and functional capabilities of the MPS, our iRECs and iRPCs can advance iBRB research and precision medicine, accelerating therapeutic discovery.

Our study is the first to derive retinal-specific ECs from iPSCs by harnessing the Norrin/Fz4 axis, a breakthrough that provides a new approach for studying iBRB development, retinal microvascular diseases, and therapeutic interventions. We anticipate that this iREC differentiation approach will advance cell therapy and disease modeling, accelerating the discovery of treatments for retinal microvascular diseases. Harnessing our protocol, patient-specific isogenic iRECs can be developed to promote iBRB regeneration and revascularization, marking a notable step forward in precision medicine.

### Limitations of the study

A significant challenge still exists in identifying and defining tissue-specific markers in ECs and pericytes. The purification of ECs from human tissue and the loss of primary cell identity during *ex vivo* culture hinder the discovery of tissue-specific markers. Moreover, the limited genotypic studies involving human RECs has led to a minimal understanding of unique biomarkers specific to RECs. As a result, while we demonstrate the genotypic and phenotypic integrity of our iRECs, studies investigating the complexities governing human REC specification and maturation are required to uncover additional biomarkers. This will enhance our ability to validate the iRECs and confirm their phenotypic and genotypic integrity. The plasticity role of pericytes makes it difficult to characterize them using only one or two exclusive markers. In this study, we carefully selected the markers that best represented retinal specificity as published in mice and compared them to primary HRPCs. However, new markers and pathways may be identified as research progress, which could be further used to characterize and generate RECs and RPCs with greater precision. An additional challenge comes with comparing published iPSC-derived brain ECs datasets with our datasets as there is a potential for batch effects. Nevertheless, our results demonstrate, for the first time, the transcriptomic differences between ECs exhibiting brain-specific and retina-specific traits.

There are also some limitations associated with our models. While the iBRB-on-a-chip is perfusable and functional, dextran perfuses through the collagen type-I hydrogel at a significant rate compared to perfusion across the cell-cell contacts within iREC networks. Consequently, we can not accurately calculate permeability across the iREC networks in 3D utilizing this system. Advancements will need to be made with the iBRB MPS to enable permeability studies in the future. However, the iBRB MPS is still very advantageous, as it can be harnessed to investigate iBRB development and maturation, study retinal microvascular diseases, and test therapeutics. Our current system also only includes iRECs and iRPCs. Incorporating other essential cell types, such as retinal astrocytes, into the iBRB MPS could further improve the biomimicry of the model. Within our OIR model, the PBS control eyes contained iRECs, displaying iREC migration and infiltration from the contralateral eye. While this may hinder some of the data analyses and statistical comparisons, it demonstrates the immunocompatibility of our iRECs and thus their therapeutic potential and translatability.

## Acknowledgments

We thank Dr. Ou for assisting with RNASeq analysis. We thank Dimitris Ntekoumes and Emma Villares for their guidance with the network analysis protocol. We thank Dr. Peter Searson and Tracy Chung from Johns Hopkins University for generously providing the iPSC-derived brain EC RNA-Seq data used in this study. Y.L. was partially funded by the Taiwan–Whiting School of Engineering/Johns Hopkins University Fellowships program, P.E. is supported by the National Science Foundation Graduate Research Fellowship Program (NSF GRFP), and E.W. is supported by the Department of Defense (DoD) through the National Defense Science & Engineering Graduate (NDSEG) Fellowship Program. We acknowledge support from the Duke Cancer Institute as part of the P30 Cancer Center Support Grant (Grant ID: P30 CA014236). This work was supported by SNT0101 from the Translational Research Institute through NASA Cooperative Agreement NNX16AO69A and EY035853 (both to S.G.).

## Materials and methods

### Maintenance and differentiation of hiPSCs

Mono-allelic mEGFP-tagged TJP1 WTC (WTC-TJP1) hiPSCs (Coriell Institute for Medical Research) or C1-2^55^ hiPSCs were used to obtain iECs and iRECs. The differentiation protocol was based on an established endothelial cell differentiation protocol with modifications^48^. Cells were cultured on a Matrigel-coated plate in mTeSR1 (STEMCELL Technologies) with the media changed daily for 3–5 days. Endothelial differentiation started with mesodermal induction when cells were 60-80% confluent. To do this, 6 µM CHIR99021 (STEMCELL Technologies) was added to the essential 6 medium (Thermo Fisher Scientific) and media was changed daily. On day two of the mesodermal induction, cells were detached using TrypLE Express (Thermo Fisher Scientific) and transfected with 2.5 µg ETV2 modRNA (TriLink Biotechnologies) by electroporation. For the iEC differentiation, after electroporation, cells were seeded at 86,000 cells/cm^2^ on a type-I collagen-coated plate with differentiation media containing Endothelial cell growth media (ECGM; Promocell). The media was supplemented with 10 µM SB-431542 (Cayman Chemical Company), 50 ng/ml VEGF-A (PeproTech), 10 µM Y-27632 (Selleckchem), and 50 ng/ml FGF-2/bFGF (PeproTech). For the iREC differentiation, after electroporation, cells were seeded at 86,000 cells/cm^2^ on a type-IV collagen-coated plate. The same differentiation media cocktail was utilized, but was additionally supplemented with 80 ng/ml Human Norrin Recombinant Protein (Boster Bio) and 5 µg/ml vitronectin (Thermo Fisher Scientific). After 24 hours, the media was changed to remove Y-27631.

C1-2 hiPSCs^55^ were used to obtain retinal pericytes following an established protocol with modifications^151^. C1-2 hiPSCs were cultured on a vitronectin-coated plate in essential 8 medium (Thermo Fisher Scientific) and the media was changed daily for 3-5 days. Pericyte differentiation started with mesodermal induction when cells were 60-80% confluent as detailed above. On day two of the mesodermal induction, cells were detached using TrypLE Express and seeded on a type-I collagen-coated plated at 2×10^4^ cells/cm^2^ with differentiation media containing ECGM supplemented with 10 µM SB-431542, 50 ng/ml VEGF-A, 10 µM Y-27632, and 80 ng/ml Human Norrin Recombinant Protein. After 24 hours, the media was changed to remove Y-27632. After that, media was changed every other day for five days.

### Purification and expansion of iECs, iRECs, and iRPCs

For iECs and iRECs, on day five of the differentiation, CD31^+^ cells were isolated with a magnetic-activated cell sorter (MACS; Miltenyi Biotech Bergisch Gladbach). Cells were washed with dPBS (Thermo Fisher Scientific) once and dissociated using TrypLE Express. After centrifuging, cells were then resuspended in 100 µl MACS buffer containing 0.5 mM EDTA (KD Medical) and 0.5% BSA (Thermo Fisher Scientific) in dPBS. 15 µl of PE-conjugated anti-human CD31 (BD Biosciences) was then added to the MACS buffer, and cells were incubated at 4°C for 10 minutes. After incubation, cells were washed with MACS buffer twice to remove unbound primary antibodies. Next, cells were incubated with 20 µl of anti-PE microbeads (Miltenyi Biotec Bergisch Gladbach) in 80 µl of MACS buffer at 4°C for 15 minutes. Cells were washed once with MACS buffer and sorted using the MS MACS separation column (Miltenyi Biotec Bergisch Gladbach). The purified CD31^+^ iECs were plated on a type-I collage-coated plate and expanded and maintained in ECGM supplemented with 10 µM SB-431542, 50 ng/ml VEGF-A, and 10 µM Y-27632. After 24 hours, the media was changed to remove Y-27632. After that, media was changed every other day. The purified CD31^+^ pre-iRECs were plated on a type-I collagen-coated plate and expanded in ECGM supplemented with 10 µM RepSox (Selleckchem) and 10% platelet poor human serum (Millipore Sigma) for 3-5 days until confluent. The purified CD31^+^ pre-iRECs then went through a second MACS with PE-conjugated anti-human Fz4 (Miltenyi Biotec Bergisch Gladbach) as described above to obtain iRECs. iRECs were plated on a type-I collagen-coated plate and expanded and maintained in ECGM supplemented with 10 µM RepSox, 10% platelet poor human serum (Millipore Sigma), and 10 µM Y-27632. After 24 hours, the media was changed to remove Y-27632 and then media was changed every other day.

For iRPCs, on day eight of the differentiation, CD31^-^ cells were isolated with a MACS to exclude CD31^+^ cells. CD31^-^ cells then went through another MACS to obtain Fz4^+^ cells. Each MACS followed the above procedure. CD31^-^Fz4^+^ cells were plated on a type-I collagen-coated plate and expanded in Pericyte Medium (ScienCell) supplemented with 80 ng/ml Human Norrin Recombinant Protein for six days. After which, retinal pericytes were plated on a type-I collagen-coated plate and expanded and maintained in Pericyte Medium.

### Primary cell culture

HRECs (Cell systems) passages 4-6 were cultured on collagen type I-coated plate in Complete Classic Medium (Cell systems) with media change every 48 hours. HRPCs (Cell systems) passages 4-6 were cultured on collagen type I-coated plate in Complete Classic Medium (Cell systems) with media change every 48 hours.

### Flow cytometry analysis

Cells were harvested for analysis using TrypLE (Invitrogen) dissociation buffer and collected in 100 µL of 0.1% BSA. Cells were then incubated with primary antibody for 30 minutes on ice. Antibodies are detailed in **Table S1**. Cells were washed three times with 0.1% BSA and filtered through a 40-µm cell strainer. Flow analysis was conducted utilizing a BD FACSCanto flow cytometer. Following manufacturer instructions, dead cell populations were gated out with forward-side scatter plots. All analyses were conducted using FlowJo software (v10.1).

To analyze total Fz4 expression, cells were harvested using TrypLE dissociation buffer. Cells were centrifuged and fixed in 2% paraformaldehyde (PFA; Sigma-Aldrich) for 15 minutes on ice and then washed two times with 0.1% BSA. Cells were then permeabilized with 0.1% TWEEN 20 (Sigma-Aldrich) in PBS for 10 minutes at room temperature, after which cells were washed two times with 0.1% BSA. Cells were incubated with Fz4 antibody for 30 minutes on ice. Cells were washed three times with 0.1% BSA and filtered through a 40-µm cell strainer. Flow analysis was conducted on a BD FACSCanto flow cytometer. Following manufacturer instructions, dead cell populations were gated out with forward-side scatter plots. All analyses were conducted using FlowJo software (v10.1).

### Quantitative reverse transcriptase PCR gene analysis

Total RNA was extracted using TRIzol reagent (Thermo Fisher Scientific) and purified using the RNeasy Mini Kit (Qiagen). RNA quality and concentration were measured using a nanodrop spectrophotometer. Complementary DNA (cDNA) was generated using GoScript Reverse Transcriptase Random Primers kit (Promega) per the manufacturer’s protocol. The TaqMan Universal PCR Master Mix and Gene Expression Assay were used for the genes of interest. TaqMan PCR was performed using the QuantStudio 3 PCR System. The results were calculated as 2^-ΔΔCT^ obtained by comparing the cycle threshold (CT) between samples as normalized to the endogenous control gene TATA-binding protein (TBP). **Table S1** lists all primers. Each biological replicate pairing across conditions was tested in parallel to account for and minimize batch effects.

### RNA-Seq sample preparation and analysis

Total RNA was extracted using TRIzol reagent and purified using the RNeasy Mini Kit. RNA quality and concentration were checked with NanoDrop and passed quality control performed by Novogene. Four biological replicates from iECs and iRECs were analyzed. Each biological replicate pairing across conditions was differentiated and collected in parallel to account for and minimize batch effects. Libraries were prepared, sequenced and analyzed by Novogene.

To compare differential gene expression in iECs, iRECs and iPSC-derived brain ECs, previously published iPSC-derived brain EC sample data were obtained from Gene Expression Omnibus database (GSE195519). RNA-Seq reads were trimmed by Trim Galore (v 0.6.7, cutadapt v 3.5) and the cleaned reads were counted using salmon ^152^ (v 1.4.0, with parameters--libType=(IU for iBMEC samples and ISR for iEC and iREC samples) and supplying the Ensembl GRCh38 transcriptome). Bioconductor package DESeq2^153^ (v 1.34.0) was employed to analyze differential expressions (DE). The volcano plots were created by EnhancedVolcano (v 1.12.0). TPM values were quantified with Salmon^154^ (v 1.4.0) and summarized via tximport^155^ (v 1.22.0). STRING database was used for network analysis and representation^156^ (https://string-db.org/). The Venn-diagrams for expressed gene were quantified with Fragments Per Kilobase of transcript per Million mapped reads (FPKM) >1. Raw data are accessible through NCBI Gene Expression Omnibus, accession number GSE284350.

### 3D collagen hydrogel vascular network formation

Collagen gels were prepared as described previously^157^. To prepare 1 mL of collagen gel solution, 800,000 cells were resuspended in 400 µL of Medium 199 (1X) media (Thermo Fisher Scientific), 40 µL of Medium 199 (10X) media (Thermo Fisher Scientific), and 350 µL of 7.1 mg/mL Rat Tail Collagen I (Corning). The pH of the solution was adjusted by titrating with 1 M NaOH (Sigma-Aldrich) up to 10 µL. 56 µL of the mixture was added to the wells of a 96-well plate. ECGM supplemented with 50 ng/ml of VEGF-A was added after gels were polymerized at 37°C for 30 minutes. Constructs were incubated for three days to allow network formation.

Gels were fixed in 2% formaldehyde for 20 minutes followed by 3 PBS washes. Gels were incubated in 0.1% Triton-X 100 (Sigma-Aldrich) for 10 minutes and washed with PBS twice before one final 30-minute wash. Next, gels were blocked in 5% BSA solution for 1 hour at room temperature. After one PBS wash, gels were incubated overnight with the primary antibodies at 4°C. Gels were washed 3 times with PBS and then incubated with secondary antibodies, such as a conjugated phalloidin probe, for 2 hours at room temperature or overnight at 4°C. A full list of primary and secondary antibodies can be found in **Table S1**. Gels were washed 3 times with PBS, stained with DAPI for 10 minutes, washed three more times with PBS, and stored in unsupplemented PBS until imaging.

### Immunofluorescence staining and imaging

iECs, HRECs, HRPCs, iRECs, and iRPCs were fixed in 3.7% PFA for 10 minutes or in ice-cold methanol for 5 minutes. Cells fixed with PFA were permeabilized with 0.1% Triton X-100. Cells were washed and blocked in 1% BSA for one hour at room temperature or overnight at 4°C. Cells were incubated with primary antibodies overnight at 4°C. The next day, cells were washed three times with 0.1% TWEEN 20 in PBS. Cells were incubated with secondary antibodies for 1 hour at room temperature or overnight at 4°C. Primary and secondary antibodies are detailed in **Table S1**. Cells were washed three times with 0.1% TWEEN 20 in PBS followed by DAPI staining for 10 minutes at room temperature. iEC, HREC, iREC, HRPC, iRPC, iBRB-on-a-chip, and construct samples were imaged on a Nikon AX-R confocal microscope using Element software. Depictions are maximum intensity projections across the entire Z-stack except for cross-section 3D depictions and unless otherwise mentioned.

### Immunofluorescence quantification

Junctional protein localization at cell-cell contacts was quantified in Fiji^158^ utilizing 60x magnification confocal images. To ensure only the protein expression at the junctions was quantified, the fluorescence on the interior of each quantified cell was removed. A cell of interest was then manually highlighted along its cell-cell contact junctions and then the mean fluorescence intensity (MFI) was measured with Fiji’s MFI measurement feature. To prevent inaccurate calculations, only clear cell-cell boundaries were quantified and background MFI was measured and subtracted from each recorded junctional MFI value.

For all *in vitro* vascular network quantification, vascular branch length, surface area, and volume quantifications were made utilizing 20x magnification confocal images while the higher resolution 40x magnification was used to calculate network lumen diameter quantification. We used Imaris (v9.6) to quantify the branch length and vascular volume across the entire z-stack of each sample. Harnessing thresholding and surface masking, we developed a filament reconstruction of each sample’s entire vascular network. Performing a subsequent filament analysis established the average vascular branch length and volume. To accurately measure vascular surface area, we utilized Fiji following a well-established protocol^158,159^. Briefly, we performed thresholding to establish the networks and then utilized a Trainable Weka Segmentation 3D module to identify the vasculature. The 3D geometrical measure plug-in was then run to quantify surface area. The vascular network lumen diameters were manually quantified via NIS-Elements AR (v6.02.03) software with the manual measurement feature. The lateral and transverse diameter was measured and averaged for each lumen to account for the non-uniform nature of lumens, following previous reported and validated work^89^. For all *in vitro* quantifications, the phalloidin channel was utilized.

For *in vivo* vascular network quantification, 10x magnification confocal images were used for all network quantification to visualize for the entire retina. Imaris (v10.0) was used to quantify branch length and network segment volume following the filament tracer procedure outlined above. Segment refers to quantifying individual portions that compose each filament. Surface area and lumen diameters were calculated in Fiji following the well-established protocol detailed above^158,159^. Lumen diameters were not manually calculated due to the significant number of networks present and low resolution of the images. For all *in vivo* quantifications, the Isolectin B4 channel was utilized.

For all quantification experiments, each biological replicate pairing across conditions was tested in parallel to account for and minimize batch effects.

### Diabetic retinopathy (hypoxia and high glucose) treatment

To recapitulate diabetic retinopathy, we treated iRECs with diabetic media, composed of ECGM supplemented with 10 µM SB-431542, 10 µM RepSox, 10% platelet poor human serum, and 30 mM D-Glucose (Sigma-Aldrich), and exposed them to 5% CO_2_, 95% N_2_ conditions (BioSpherix), resembling hypoxia. For the junctional protein localization and RT-qPCR analyses, a 24-hour diabetic treatment was performed when iRECs reached a confluency of ∼80 – 90%. The iRECs undergoing TEER assays were exposed to a 72-hour diabetic treatment beginning on day 1. Diabetic media was changed daily. For the iREC vascular networks developed in collagen-I hydrogels, we provided a 24-hour diabetic treatment to the networks on day 2. The control for each experiment included iRECs exposed to 25% O_2_ conditions and treated with ECGM supplemented with 10 µM SB-431542, 10 µM RepSox, and 10% platelet poor human serum. Sample collection was performed at the conclusion of diabetic treatment for all experiments.

### Transendothelial electrical resistance (TEER) assay

The transendothelial electrical resistance (TEER) assay was performed following a standard, well-established protocol^146^. In short, iECs and iRECs were plated (day 0) on 24-well plate Transwell inserts containing 0.4 µm pores (Corning). For the iREC vs. iEC testing, media was changed on day 1 and day 3 and TEER values were measured 24 ± 2 hours apart between day 1 and day 4. For the DR recapitulation experiments, media was changed daily to avoid dehydration and TEER values were measured 24 ± 2 hours apart between day 1 and day 4. The first measurement was made one hour after treatment induction, acting as an internal control that ensured iREC consistency and accuracy prior to experiencing sufficient diabetic conditioning to induce barrier property alteration. All TEER measurements were made using the EVOM3 system and an STX-4 probe (World Precision Instruments). To calculate the reported Ω cm^2^ TEER values, the recorded resistance was subtracted from the resistance of just the media within the same Transwell insert and then corrected for the Transwell membrane area (0.33 cm^2^). Each biological replicate pairing across conditions was tested in parallel to account for and minimize batch effects.

### GLUT1 uptake assay

The Glucose Uptake-Glo^TM^ Assay kit (Promega, J1341) was implemented to perform the GLUT1 uptake assay. iRECs were cultured on 6-well tissue culture plates (Corning) until they reached 80 – 90% confluency. Three wells were then treated with 2 µM of the highly selective GLUT1 inhibitor, Bay-876^160–162^ (Sigma-Aldrich), overnight before the assay and during the experiment. Three wells were left untreated as the control. At the end of the overnight treatment, all six wells were rinsed twice with PBS followed by the addition of 1000 µL of PBS supplemented with 1 mM 2-deoxyglucose (2DG) and a 10 min incubation at room temperature. For the wells treated with BAY-876, their PBS-2DG solution was supplemented with 2 µM BAY-876 prior to the 10-minute incubation period. Each well was then treated with 500 µL of Stop Buffer.Next, 75 µL of each well’s supernatant was transferred to a 96-well plate and 25 µL of Neutralization Buffer was added to each well as along with 100 µL of 2DG-6-phosphate Detection Reagent. The plate was then incubated for one hour at room temperature. A standard plate reader was utilized to measure luminescence via a 0.3 second integration. To account for background luminescence, each set of experiments was accompanied by a blank supernatant (2DG solution unexposed to the detection reagent) that was transferred to the 96-well plate.

### Oxygen induced retinopathy mouse model and intravitreal injection

We followed a well-established OIR mouse model protocol^44^. C57BL/6J (Jackson Laboratory) mice were used for the experiments. Mice were subjected to 75% oxygen from P7 to P12 (BioSpherix ProOx 360). On P12, female or male mice were intravitreally injected with PBS as vehicle control in one eye and iRECs diluted in PBS (50,000 cells/µl) in the contralateral eye by using a microinjector. All studies were conducted under the Duke University Institutional Animal Care and Use Committee-approved animal protocol A070-22-04.

### Retina flat mount and immunofluorescence staining

Eyes from P17 pups were enucleated and immediately fixed in 4% PFA for 30 minutes. The cornea, leans, and retina were removed from the eye cup using dissection forceps and micro scissors under a dissecting scope. The isolated retinas were blocked for 1.5 hours using 10% normal goat serum (Vector Laboratories) containing 0.3% Triton X-100 in PBS. The retinas were then incubated with DyLight 649-conjugated Griffonia Simplicifolia Lectin I (GSL I) Isolectin B4 (Vector Laboratories) and Rhodamine-conjugated Ulex Europaeus Agglutinin (UEA 1, Vector Laboratories) overnight at 4°C in 5% normal goat serum containing 0.3% Triton X-100 in PBS. Retinas were washed and mounted on the slide using Fluoromount-G (Invitrogen). Images of the samples were obtained using a Nikon AX-R Confocal microscope using Element software. Avascular area and neovascular tufts were quantified following a well-established protocol^82^. Briefly, we compared the number of pixels in the area of VO (VO%) or neovascular tufts (NV%) with the total number of pixels of the retina. Mice with body weight lower than 4 g on the day of tissue harvest (P17) were excluded from analysis.

### Fabrication of microfluidic device and iBRB-on-a-chip network formation

We utilized a standard soft lithography technique to fabricate iBRB-on-a-chip MPS containing three cell channels^109^. In short, a photomask containing devices with three cell-gel channels, each 800 μm wide × 1,300 μm long × 250 μm high, was coated with PDMS, cured at 80 °C overnight, and cut out to create the template molds. The devices were then bonded to glass slides via ozone treatment (Harrick Plasma) and cured for one minute at 80 °C. To sterilize the MPS templates, both sides of the devices were exposed to UV-light for 30 minutes and then the channels were coated with Poly-L-Lysine (Sigma-Aldrich) for one hour, rinsed three times with distilled water, and coated with distilled water and left on a rocker overnight. The device was rinsed one final time with distilled water and both sides of the MPS templates were exposed to UV light for 30 minutes.

To develop vascular networks we followed a published protocol^109,159^. Cells were resuspended in 12.59 µL/gel of ECGM supplemented with 50 ng/ml of VEGF-A at 340,000 iRECs/gel. They were then dispensed into a hydrogel solution containing 13.13 µL/gel of M199 1X media, 4.54 µL/gel of ECGM supplemented with with 50 ng/ml of VEGF-A, 2.4 µL/gel of M199 10X media, 27.34 µL/gel of Collagen-I, and 2.50 µL/gel of Neutralizing Solution (Advanced Biomatrix). 10 µL of the solution was loaded into each MPS cell channel and cured for 30 minutes at 37 °C. The sink and source ports within each device received 100 µL of ECGM supplemented with 50 ng/ml of VEGF-A on day 0. Media ports were changed daily until day 3. For the iREC-iRPC co-culture iBRB-on-a-chip, we resuspended the iRECs and iRPCs in 12.59 µL/gel of ECGM supplemented with with 50 ng/ml of VEGF-A at a 15:1 iREC:iRPC ratio, maintaining 340,000 iRECs/gel. We then followed the aforementioned MPS network fabrication process to develop iREC networks encapsulated by iRPCs. Both the mono- and co-culture devices were fixed and processed on day 3 for analysis.

Devices were fixed in 4% formaldehyde for 20 minutes followed by 3 PBS washes. Gels were then incubated in 1% Triton-X 100 for 20 minutes and washed with PBS twice. Next, gels were blocked in 5% BSA solution for 1 hour at room temperature. After one more PBS wash, gels were incubated overnight with the primary antibodies at 4°C. Gels were then washed 3 times with PBS containing 0.05% TWEEN 20 and incubated with secondary antibodies for 2 hours at room temperature or overnight at 4°C. A full list of primary and secondary antibodies can be found in **Table S1**. Gels were washed 3 times with PBS containing 0.05% TWEEN 20, incubated with DAPI for 10 minutes at room temperature, washed three times with non-supplemented PBS, and stored in non-supplemented PBS until imaging.

### Statistics

Biological replicates (indicated as N) are defined as separate differentiation replicates and technical replicates (indicated as n) come from the same N cultured in different tissue cultures well. All *in vitro* quantifications are N = 3 with n = 3 unless otherwise indicated in the figure legends. For *in vivo* experiments, mice number in each experiment is indicated as n. All bar graphs represent means ± SD.

Graphpad Prism (v10.2.3.) was utilized for statistical analysis. For comparisons across physiological conditions, such as iECs vs. iRECs, two-tailed unpaired Student t-tests were employed, while comparisons between physiological and diabetic conditions implemented two-tailed paired Student t-tests. Significance levels were set at *p* > 0.05, *\*p* ≤ 0.05, *\*\*p* ≤ 0.01, *\*\*\*p* ≤ 0.001 and *\*\*\*\*p* ≤ 0.0001.

## Notes

### Competing Interest Statement

The authors have declared no competing interest.

