## Supplementary material for "Derivation of functional retinal endothelial cells from human pluripotent stem cells for therapeutics and modeling"

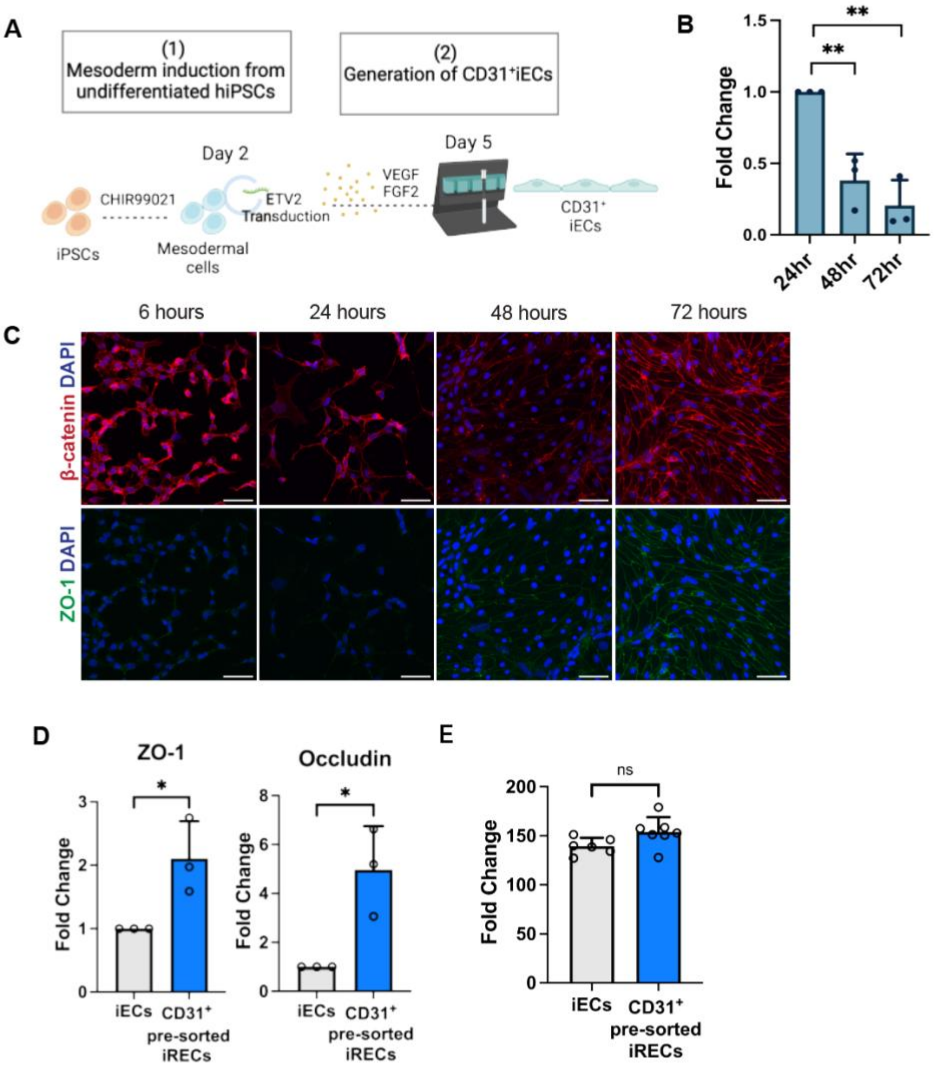

**Figure S1. Generation of CD31$^{+}$ iECs and pre-iRECs from hiPSCs and functional analysis.** (A) Schematic illustration of iEC differentiation. (B) RT-qPCR of ETV2 expression for pre-iRECs. (C) Representative immunofluorescence staining of adherens and tight junction proteins along the pre-iREC differentiation. β-catenin (red), ZO-1 (green), and nuclei (blue). Scale bars: 50 µm. (D) RT-qPCR for iECs and CD31$^{+}$ pre-sorted iRECs on day 8. (E) TEER measurements of iECs and CD31$^{+}$ presortediRECs. N = 3, n = 2 for each sample for TEER measurement.

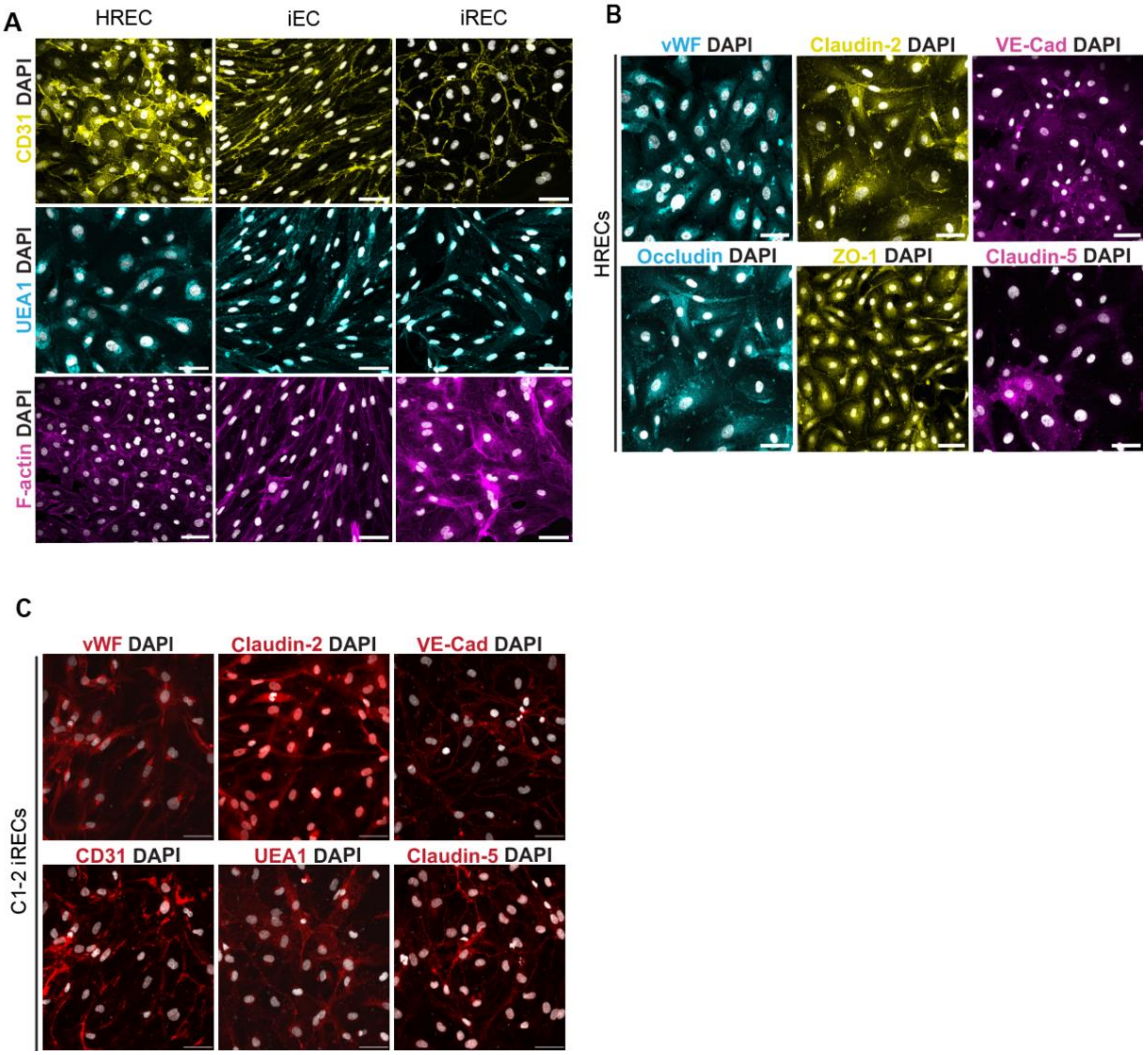

**Figure S2. Immunofluorescence images of HRECs, iECs, and iRECs.**
(A) Representative immunofluorescence images of EC markers in HRECs, iECs, and IRECs. CD31 (yellow), UEA1 (cyan), F-actin (magenta), and nuclei (white). (B) representative immunofluorescence images of EC and junctional protein markers in HRECs. Claudin-5, and VE-Cadherin (magenta), Claudin-2 and ZO-1 (yellow), vWF and Occludin (Cyan), and nuclei (white). (C) representative immunofluorescence images of EC and junctional protein markers in C1-2 hiPSC-derived iRECs. vWF, Claudin-2, VE-Cad, CD31, UEA1, and Claudin-5 (red) and nuclei (white). Scale bars: 50 µm
.

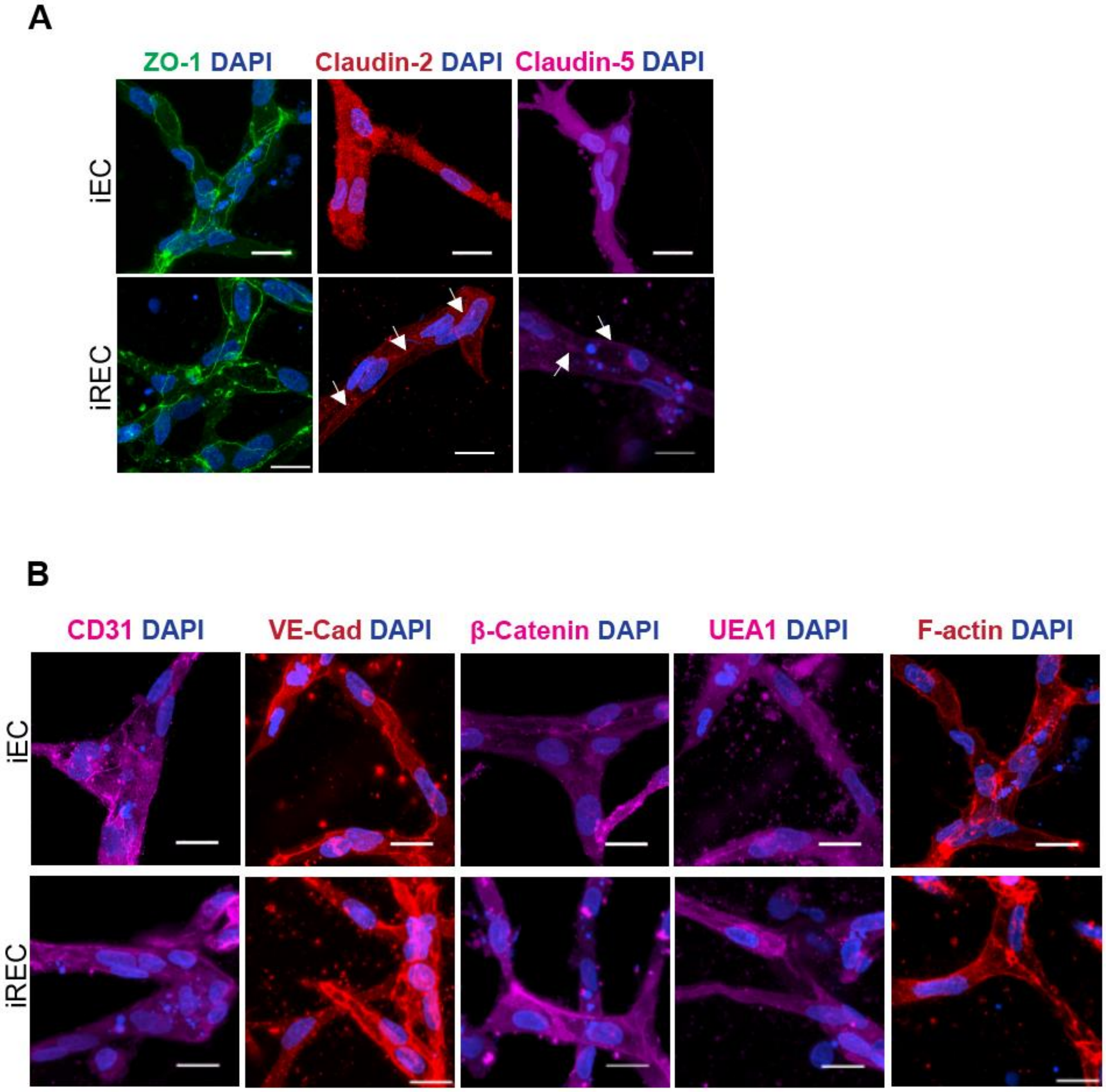

**Figure S3. Immunofluorescent images of 3D iEC and iREC vascular networks.**
Representative immunofluorescence images of (A) junctional protein expression at cell-cell contacts (some indicated with arrows) and (B) endothelial cell marker expression in 3D iEC and iREC vascular networks. ZO-1 (green), Claudin-2, VE-Cadherin, and F-actin (red), Claudin-5, CD31, β-catenin, and UEA1 (magenta), and nuclei (blue). Scale bars: 20 µm.

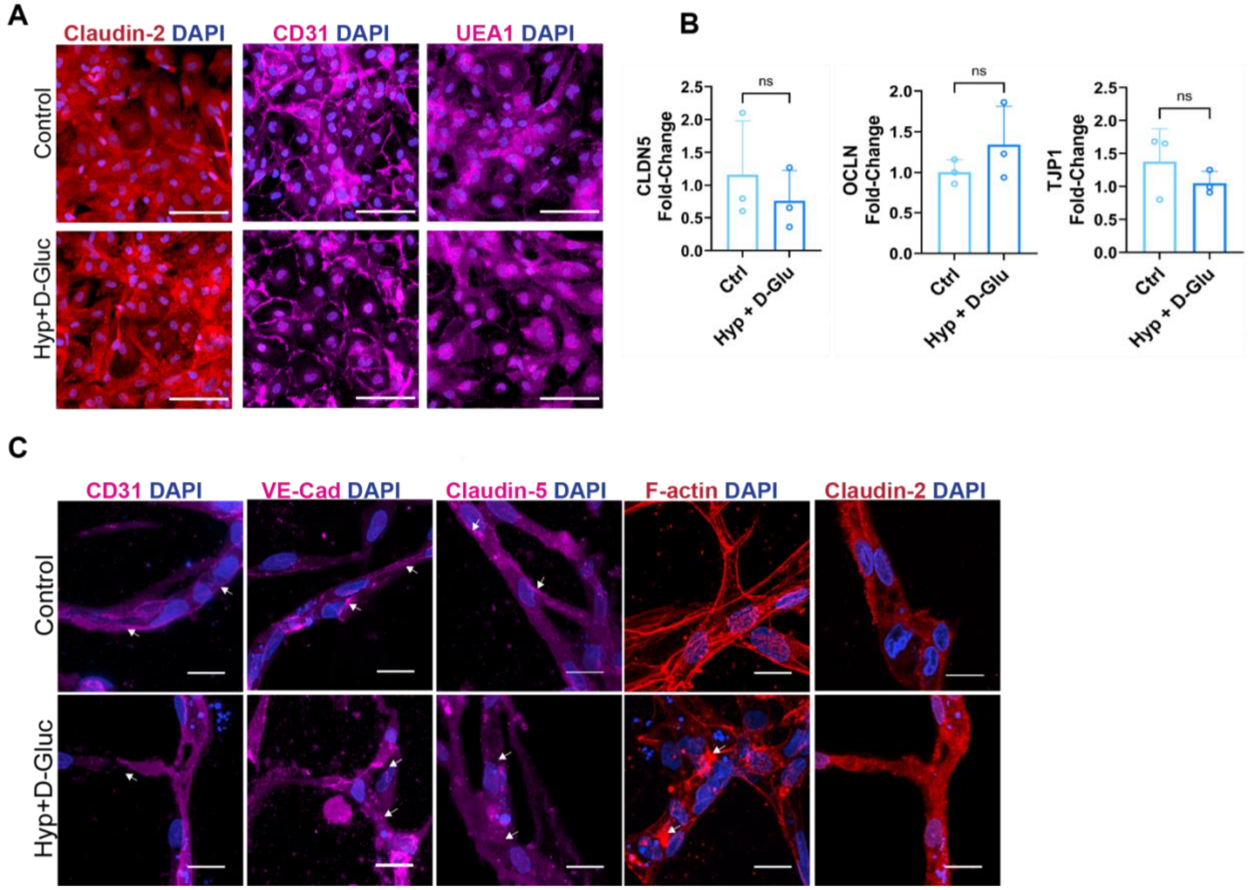

**Figure S4. iREC phenotype in healthy and diabetic conditions.**
(A) Representative immunofluorescence images of iRECs with or without diabetic treatment (Hyp + D-Gluc). Claudin-2 (red), CD31 and UEA1 (magenta), and nuclei (blue). (B) RT-qPCR analysis for Claudin-5 (CLDN5), Occludin (OCLN), and ZO-1 (TJP1) within iRECs exposed to diabetic or control conditions. (C) Representative immunofluorescence images of 3D iREC vascular networks expressing endothelial cell markers and junctional proteins at cell-cell contacts (arrows) exposed to diabetic or control conditions. Scale bars: 100 µm (A) and 20 µm (C).

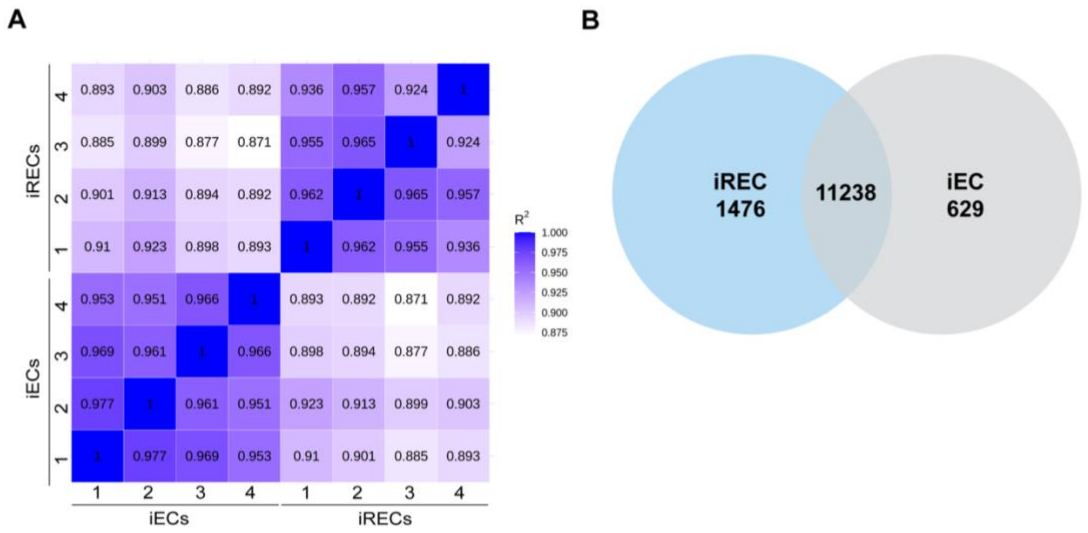

\

**Figure S5. Bulk RNA-Seq analysis of iECs and iRECs.**

(A) Correlation analysis and (B) Venn diagram illustrating the differential gene expression between iECs and iRECs across four biological replicates

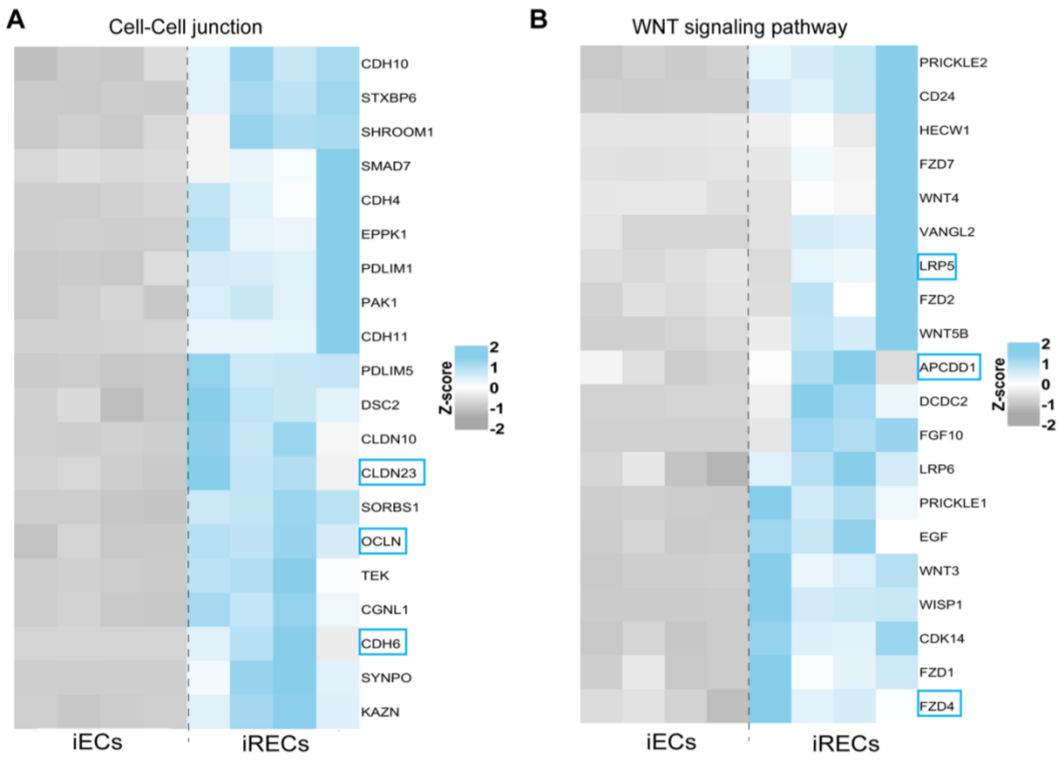

**Figure S6. Bulk RNA-Seq analysis of iECs and iRECs.**
Heatmap analysis of (A) cell-cell junction-related genes and (B) Wnt signaling pathway-related genes in iECs and iRECs across four biological replicates.

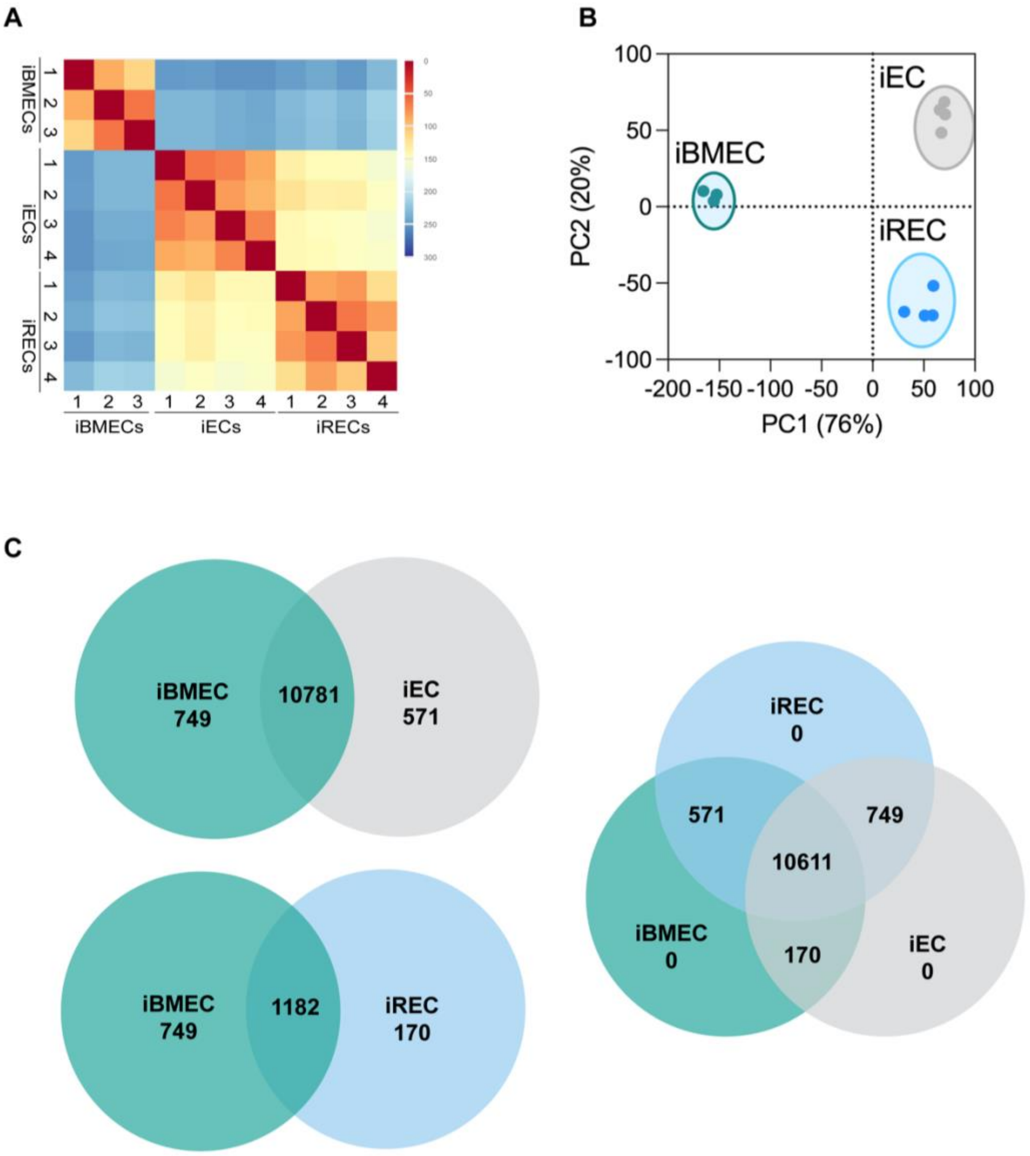

**Figure S7. Bulk RNA-Seq analysis of iECs, iRECs and iPSC-derived brain ECs.**
(A) Correlation analysis of iECs, iRECs and iPSC-derived brain ECs across four biological replicates. (B) Principal component analysis of RNA-Seq data from iECs, iRECs and iPSC-derived brain ECs. (C) Venn diagram illustrating the differential gene expression between iPSC-derived brain ECs, iECs and iRECs.

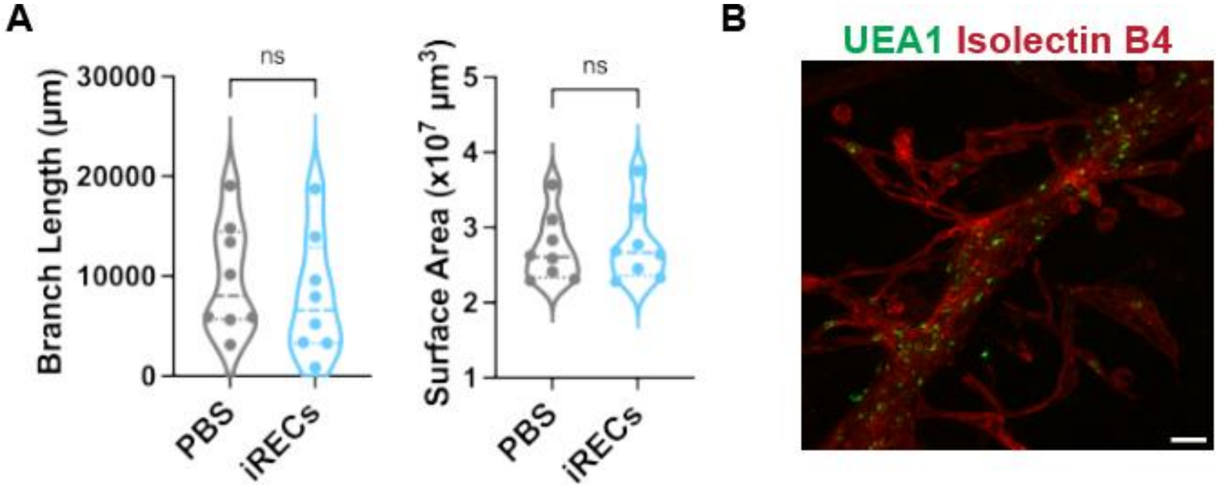

**Fig S8. PBS and iREC intravitreal injections into OIR mouse model.**
(A) Quantification of mouse vascular network branch length and surface area on postnatal day 17 (P17) in PBS and iREC injected groups. (B) An immunofluorescence image of iRECs along the mouse retinal artery on P17 within a PBS treated eye. Mouse isolectin B4 (red) and human UEA1 (green). Scale bars: 25 µm.

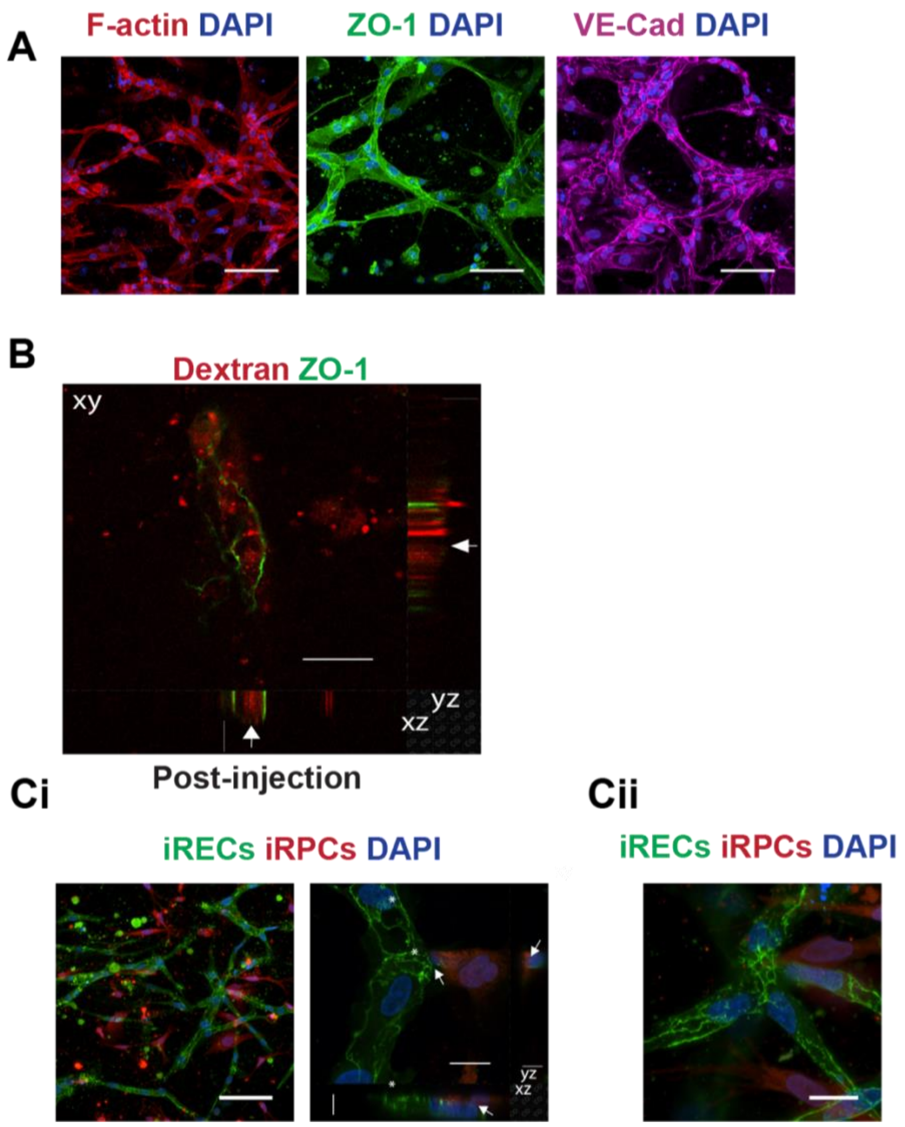

**Figure S9. Characterization of iBRB-on-a-chip.**
(A) Representative immunofluorescence images of 3D iREC vascular networks expressing tight and adherens junction proteins at cell-cell contacts. F-actin (red), ZO-1 (green), VE-Cadherin (magenta), and nuclei (blue). (B) Cross-section of 70 kDa rhodamine dextran perfusing iREC networks following the perfusion assay. ZO1 (green) and 70 kDa rhodamine dextran (red). Arrows denote dextran perfusion through the lumen of an iREC network. (C) Representative immunofluorescence images of the iREC-iRPC co-cultured iBRB-on-chip displaying perivascular iRPCs (red) surrounding and encapsulating iREC networks (green) in order to support their functional lumens. Asterisks denote lumens and arrows highlight iRPC-iREC interactions. Scale bars: 100 µm (A and Ci left panel), 20 µm (B, Ci xy plane in the right panel, and Cii), and 10 µm (Ci xz and yz planes in the right panel).

**Table S1. Key Resources.**

| | | | | | |
|---|---|---|---|---|---|
| Calponin-1 | SCBT | sc-58707 | ICC | Mouse | 1/100 |
| PDGFRb | SCBT | sc-432 | ICC | Rabbit | 1/200 |
| CD13 | SCBT | sc-18899 | ICC | Mouse | 1/100 |
| Nestin | SCBT | NB300-265s | ICC | Rabbit | 1/200 |
| NG2 | SCBT | sc-166251 | ICC | Mouse | 1/100 |
| Transgelin (SM22α) | Cell Signaling | 40471s | ICC | Rabbit | 1/100 |
| CD31 | Abcam | ab28364 | ICC | Rabbit | 1/100 |
| VE-Cadherin | SCBT | sc-9989 | ICC | Mouse | 1/100 or 1/4000 |
| von Willebrand Factor | SCBT | sc-365712 | ICC | Mouse | 1/100 |
| Alexa Fluor 546 Phalloidin | Thermo Fisher | A22283 | ICC | N/A | 1/400 |
| Occludin | Thermo Fisher Scientific | 711500 | ICC | Rabbit | 1/100 |
| Claudin-5 | Thermo Fisher Scientific | 35-250-0 | ICC | Mouse | 1/50; 1/200 |
| ZO-1 | Thermo Fisher Scientific | 40-2200 | ICC | Rabbit | 1/100 |
| Claudin-2 | Thermo Fisher Scientific | 710221; 5600 | ICC | Rabbit; Mouse | 1/200; 1/100 |
| Dylight-conjugated Ulex Europaeus Agglutinin | Vector Laboratories | DL-1068-1 | ICC | N/A | 1/400 |
| DAPI | Thermo Fisher Scientific | D1306 | ICC | N/A | 1/1000 |
| ***In vivo*** | | | | | |
| Griffonia Simplicifolia Lectin I (GSL I) Isolectin B4, DyLight™ 594 | Vector Laboratories | DL-1207-.5 | IF | N/A | 1/400 |
| Anti-NG2, Alexa Fluor™488 Conjugate Antibody | Millipore Sigma | AB5320A4 | IF | Rabbit | 1/200 |
| Rhodamine-conjugated Ulex Europaeus Agglutinin | Vector Laboratories | RL-1062-2 | IF | N/A | 1/400 |
| **Flow cytometry** | | | | | |
| PE-Anti-Human CD140b | BD Pharmingen | 558821 | Flow cytometry | Mouse | 1/10 |
| APC-Anti-Human CD344 | Miltenyi Biotec | 130-106-614 | Flow cytometry | Mouse | 1/10 |
| PE-Anti-Human CD31 | BD Pharmingen | 555446 | Flow cytometry | Mouse | 1/10 |
| **Primers** | **Taqman assay** | | | | |
| Claudin-5 | Hs00533949_s1 | | | | |
| Lrp5 | Hs00182031_m1 | | | | |

| | |
|---|---|
| ZO-1 | Hs01551861_m1 |
| Occludin | Hs00170162_m1 |
| TBP | Hs00427620_m1 |
